# High frame rate *in vivo* two-photon microscopy to quantify murine cerebrospinal fluid flow heterogeneity

**DOI:** 10.64898/2026.09.04.749497

**Authors:** Cooper Gray, Kaidi Hu, Dorothea H. Tse, Kyle Dieterle, Silas Simpson, Thomas Ruhl, Kyle Duff, Daehyun Kim, Mahsa Mirzaee, Anika Volker, Turki S. Alturki, Jeffrey Tithof

## Abstract

Cerebrospinal fluid (CSF) flows through perivascular spaces (PVSs) surrounding brain vasculature, and impaired flow has been linked to neurodegenerative diseases such as Alzheimer’s. However, the mechanisms driving CSF flow and its oscillatory dynamics remain poorly understood. Using high frame rate two-photon imaging (up to 113 Hz), we show that CSF flow is spatially heterogeneous and highly pulsatile, quantified using a regional CSF pulsatility index (PI). Reduced-order simulations with realistic domain length and Windkessel boundary conditions show that arterial pulsations generate peak CSF velocities but contribute negligibly to net transport. PI varies non-monotonically in space and substantially with downstream hydraulic resistance and compliance, indicating that CSF pulsatility is an emergent property of the hydraulic network rather than simply a consequence of local arterial wall motion. Measurements and simulations of the phase between arterial wall motion and CSF velocity further constrain proposed CSF driving mechanisms. Finally, we perform mouse-specific simulations to estimate average and peak wall shear stresses which support the potential role of pulsatile CSF flow in perivascular mechanotransduction. Together, these results establish CSF pulsatility as a measurable signature of network-scale CSF dynamics.

---

The past decade has seen a rapid increase in scientific interest in characterizing cerebrospinal fluid (CSF) circulation through the glymphatic (glial-lymphatic) system [1, 2]. This pathway involves bulk CSF flow through perivascular spaces (PVSs), annular channels surrounding all blood vessels in the brain that facilitate solute exchange with interstitial fluid. While early studies focused on glymphatic function in mice [3, 4, 5], a growing body of literature suggests humans possess a glymphatic system with many of the major features reported in rodents [6, 7, 8]. Impaired glymphatic function has been associated with neurodegenerative disease, stroke, and traumatic brain injury, motivating efforts to understand the mechanisms governing CSF transport [9, 10, 11, 12].

A rapidly growing body of research suggests multiple mechanisms drive CSF through the glymphatic system under physiological conditions. Cardiac arterial pulsations have substantial experimental support [13, 14, 4, 15, 16, 5], but their small amplitude (about 1% the vessel diameter [5]) and long wavelength, longer than the entirety of the vasculature, have raised questions about their ability to generate observed net flows in a variety of numerical studies [17, 18, 19, 20]. Respiration also modulates CSF transport through autonomic modulation of cardiovascular function [21] and alterations in intracranial pressure [22]. Emerging evidence also points to slow vasomotion [23] and neurovascular coupling during non-rapid eye movement (NREM) sleep as major drivers [7, 24], which generate arterial pulsations with amplitude as large as about 10%, but at frequencies *>* 100 times lower than cardiac pulsations. Even if these latter mechanisms dominate CSF transport during NREM sleep, there are many important open questions regarding the role of cardiac pulsations during other states, including rapid eye movement sleep, anesthesia [5], or pathologies in which vasomotion is diminished [25].

CSF flow in PVSs at the surface of the mouse brain has been directly quantified in a handful of prior experimental studies in which fluorescent microspheres were injected into the CSF then imaged at high spatial resolution [5, 26, 27]. However, these studies used a relatively slow frame rate (30 Hz), which yielded about five measurements per cardiac cycle (an anesthetized mouse has a heart rate of about 6 Hz), which is inadequate for fully resolving the temporal fluctuations in flow velocity and associated quantities such as the peak wall shear stress (WSS). Accurate measurements of WSS are essential for evaluating the potential role of mechanotransduction in PVSs, which has been shown to regulate cardiovascular [28] and lymphatic [29] formation. Adequately resolved flow measurements are also essential for calibrating numerical models that test hypotheses about mechanisms of glymphatic transport [18, 19, 20, 30]. Such prior numerical simulations also have major limitations: they often assume unrealistically short domains in which cardiac pulsations necessarily drive miniscule net flows, and they impose unrealistic boundary conditions that do not accurately reflect the highly resistive and compliant channels further downstream [18, 31, 32].

Results presented below quantify *in vivo* CSF flow through murine pial PVSs at frame rates up to 113 Hz, enabling characterization of peak CSF velocities and a CSF pulsatility index (PI). We use reduced-order finite volume numerical simulations with realistic Windkessel boundary conditions to assess sources of pulsatility variation in experiments; we find that flow pulsatility is considerably sensitive to downstream hydraulic resistance, which suggests that measurements at the surface of the brain may be used to probe properties of deeper brain flow pathways. We also characterize the phase offset between arterial pulsations and CSF velocity in experiment and simulation, which provides quantitative insight into the feasibility of recently-proposed CSF driving mechanisms. Finally, we present mouse-specific finite element simulations to estimate peak and mean WSS in PVSs. These results suggest that flow pulsatility likely plays an important role in mechanotransduction, providing a potential mechanism by which CSF flow may influence perivascular cell function and regulate glymphatic homeostasis.

## Results

### Quantitative measurements of flow heterogeneity

We anesthetized mice with ketamine-xylazine (KX), which largely preserves glymphatic transport compared to other anesthetics like isoflurane [33, 34], then implanted a cranial window over the dorsal branches of the middle cerebral artery (MCA) and injected fluorescent microspheres into the CSF. We measured CSF flow by imaging microspheres flowing in pial PVSs using two-photon microscopy (2PM), then performing particle tracking velocimetry (PTV; Fig. 1a) [35]. Reducing the field of view (FOV) along the slow (galvanometric) scanning direction increased the frame rate from the standard 30 Hz used in prior studies [5, 26, 27] to as high as 113 Hz (and 226 Hz for one mouse; see Methods). At 113 Hz, we obtain up to about 19 PTV measurements per cardiac cycle, compared to about 5 measurements at 30 Hz (assuming a heart rate of 6 Hz for an anesthetized mouse). This improved temporal resolution enables higher-fidelity measurements of the transient particle velocity peaks associated with each cardiac cycle.

**Figure 1:**
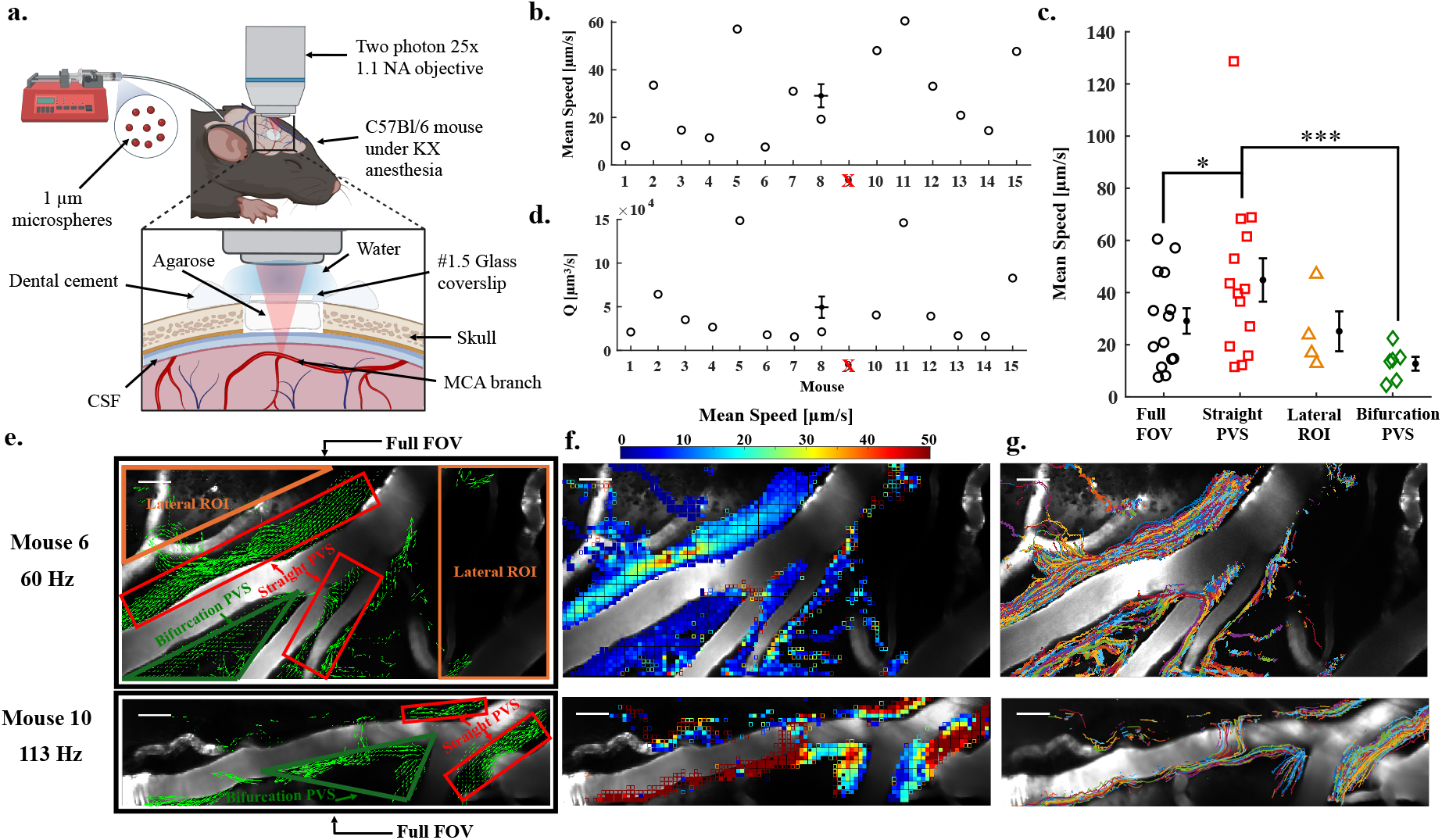
High-frame rate two-photon microscopy and particle tracking velocimetry were performed to assess CSF transport across multiple regions of interest (ROIs). **(a)** CSF flows were measured using fluorescent microspheres delivered via an implanted cannula in the cisterna magna and imaged through a glass cranial window in an anesthetized mouse. Microspheres were imaged flowing along PVSs of pial arteries; vasculature was visualized using FITC dextran delivered to blood via retro-orbital injection. **(b)** Mean flow speeds for the full FOV for each mouse (mouse 9 had a mean flow speed of 227 µm/s, which was excluded as an outlier; see Fig. S2 for separate analysis). The error bar near the middle indicates the mean ± SEM for *n* = 14 mice. **(c)** Mean speeds were calculated for specific ROIs. Statistical comparisons were performed using a linear mixed-effects model with ROI and frame rate as fixed effects and mouse and recording as random intercepts, followed by pairwise ROI comparisons using Holm correction across the six possible ROI comparisons. Mean speed was greater in Straight PVSs than Full FOV ROIs (Holm-adjusted *p* = 0.0107) and Bifurcation PVS ROIs (Holm-adjusted *p* = 2.10 × 10^−4^). Error bars indicate mouse-level mean ± SEM; *n* = 14 Full FOV, *n* = 14 Straight PVS, *n* = 6 Bifurcation PVS, and *n* = 4 Lateral ROI mice. \**p <* 0.05, \*\**p <* 0.01, \*\*\**p <* 0.001. **(d)** Mean volumetric flow rates *Q* were calculated for each mouse by estimating PVS cross-sectional areas (based on in-plane PVS width) and multiplying by the mean speed of the straight PVS ROI. The error bar near the middle indicates the mean ± SEM, *n* = 14 mice. **(e)** Time-averaged velocity fields for 60 Hz and 113 Hz show downstream direction of transport in straight PVSs, some reverse flow in the bifurcation PVS, and flow in lateral ROIs far away from the vessel. **(f)** Time-averaged flow speeds show parabolic-like flow in the straight PVSs, as well as spatially heterogeneous speeds depending on the ROI. **(g)** PVS widths were estimated in straight PVSs by superimposing all recorded tracks. Scale bars: 40 µm.

Our PTV analysis includes 15 mice with an average of 12,000 particles tracked per recording (see Source Data), which yielded a mean speed (spatially averaged over the full FOV for each mouse) of 29.1 ± 4.9 µm/s (Fig. 1b). We also quantified flow in anatomically different regions of interest (ROIs) within the FOV which revealed heterogeneity in the mean speed (Fig. 1c). This finding likely explains why our mean flow speed measurements are larger than prior studies. For example, Mestre et al [5] primarily performed PTV measurements in regions that included bifurcation PVS ROIs. We measured mean flow speeds of 12.7 ± 2.7 µm/s in bifurcation PVS ROIs, compared to 44. ± 8 8.3 µm/s in straight PVS ROIs. Straight PVS ROIs were present in all datasets and exhibited mean speeds approximately 3.5-fold greater than bifurcation ROIs and 1.5-fold greater than full-FOV spatial averages. Additionally, in several datasets we observed transport of tracers in the imaging plane distant from any vessel in view and moving unconfined to the PVS which we labeled as “lateral ROIs.” Particles flowing in these lateral ROIs had substantial mean flow speeds of 25.1 ± 3.3 µm/s.

We also measured arterial radius for each mouse using custom Matlab codes [35], yielding an average of 24.5 ± 1.3 µm, consistent with a prior study which reported 23 µm [5]. PVS width was quantified by superimposing particle tracks and measuring their width along the vessel (Fig. S1) yielding 28.3 ± 1.4 µm. We estimated the total, two-lobe PVS cross-sectional area based on the assumption that the PVS was shaped like an ellipse with the semi-minor axis length equal to the arterial radius *R*_1_ [36], such that 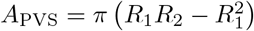, where *R*_2_ is the semi-major axis length, equal to *R*_1_ plus the PVS width (*R*_1_ and PVS width are plotted in Fig. S1c-d). The resulting PVS areas allow us to approximate PVS-to-artery area ratios (Fig. S1e). PVS and artery areas were strongly correlated (*R*^2^ = 0.75; Fig. S1f), indicating that larger arteries tended to be associated with larger surrounding PVSs. The average PVS-to-artery area ratio across our datasets was 1.16 ± 0.08, similar to previously reported values:1.4 ± 0.1 in Mestre et al. [5], 1.26 in Schain et al. [37], 1.12 in Raicevic et al. [38], and a smaller value of 0.43 reported by Smets et al. [39]. This similarity suggests that our approach provides a reasonable estimate of pial PVS geometry.

The average experimental time-averaged volume flow rate through one straight PVS lobe was 49500 ± 12000 µm^3^/s (Fig. 1d), considerably larger than previously reported values (22200 and 28500 µm^3^/s from [27, 40], respectively). These differences in *Q* arise primarily because of the larger velocities we measure in straight PVSs. One possible explanation is that the slower image acquisition used in previous studies made it more difficult to accurately resolve the fastest-moving particles, leading to lower measured velocities.

### CSF flow pulsatility index

The “pulsatility index” (PI) is a quantity routinely measured in the cardiovascular system which is defined as the difference in peak systolic and minimum diastolic velocities, normalized by the mean velocity. PI is heavily dependent on the downstream resistance in the cardiovascular network [41]. Clinicians routinely use ultrasound to noninvasively measure PI to assess vascular diseases and fetal health or even to diagnose cognitive impairment [42, 43, 44]. While numerous prior studies have investigated pulsatility of CSF flow through PVSs [4, 5, 14, 30], no prior studies have computed a PI for such flow.

First, to visualize CSF pulsatility, we plotted zoomed-in snapshots of flowing microspheres with pseudo-colored tracks indicating the particle speed over the previous several frames (Fig. 2a-b); this revealed rainbow patterns capturing velocity fluctuations over each cardiac cycle, well-aligned with unit vectors of the time-averaged velocity field (pink arrows). An animation of these tracks is included as Supplementary Video 1. We quantified the downstream component of CSF velocity, *V*_downstream_, as described in the Methods and plotted a sample time series for different regions in the FOV (Fig. 2a,c). Straight PVS ROIs tended to exhibit large positive peaks, while bifurcation and lateral ROIs had lower magnitudes and changed sign over each cardiac cycle.

**Figure 2.**
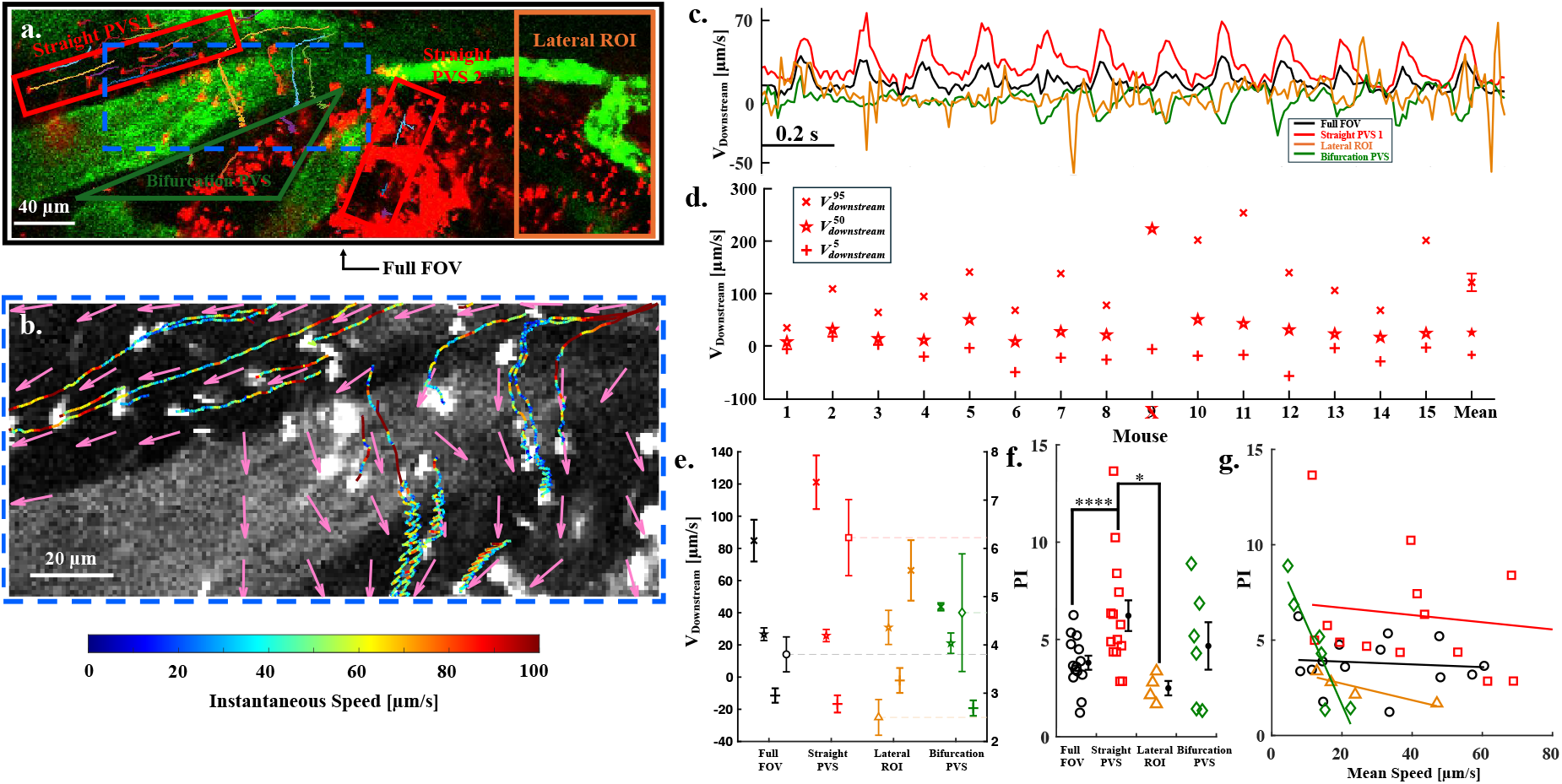
CSF flow pulsatility was quantified by computing the pulsatility index (PI) based on the maximum and minimum downstream velocity, which was adequately resolved via high frame rate imaging. **(a)** Colorful tracks show microsphere trajectories as they flow through different ROIs. **(b)** Zoomed view of blue dashed region in panel (a); rainbow tracks indicate instantaneous speed of micro-spheres over the previous frames. The rainbow patterns indicate flow pulsatility, and the pink arrows correspond to the unit vectors of the time-averaged (net) flow. See Supplementary Video 1. **(c)** Plot of two second snippet of *V*_downstream_ from (a) for each ROI. Enhanced pulsations are visible in the straight PVS relative to the full FOV, lateral ROI, and bifurcation PVS indicating that proximity to the artery enhances pulsation. **(d)** Maximum, minimum, and median values of *V*_downstream_ in the straight PVS ROI for each mouse. Error bar at the far right indicates mean ± SEM, *n* = 14 mice; the error bars for the median and minimum are smaller than the plotted symbols. **(e)** Average maximum, minimum, and median *V*_downstream_ values (left *y*-axis), and corresponding average PI values (right *y*-axis) computed across all mice for each ROI. **(f)** PI across all ROIs for all mice. Statistical comparisons used a linear mixed-effects model with ROI and frame rate as fixed effects and mouse and recording as random intercepts, followed by pairwise ROI comparisons using Holm correction across the six possible ROI comparisons. PI was greater in Straight PVS than Full FOV ROIs (Holm-adjusted *p* = 2.67 × 10^−5^) and Lateral ROIs (Holm-adjusted *p* = 0.0148). Error bars indicate mouse-level mean ± SEM; *n* = 14 Full FOV, *n* = 14 Straight PVS, *n* = 6 Bifurcation PVS, and *n* = 4 Lateral ROI mice. \**p <* 0.05, \*\**p <* 0.01, \*\*\**p <* 0.001, \*\*\*\**p <* 0.0001. **(g)** PI versus mean speed shows negative trends in all ROIs. Associations were assessed using Spearman rank correlations with Holm correction across the four ROI-specific correlations. No correlation remained statistically significant after correction, although Bifurcation PVS ROIs showed a strong negative association (*ρ* = −0.943, Holm-adjusted *p* = 0.0667).

We computed power spectra for both the arterial diameter and *V*_downstream_ which confirmed that the CSF pulsatility was locked to the cardiac frequency in 14 out of 15 cases (Fig. S3), in agreement with prior studies [5, 26, 40]. One anomalous case, mouse 9, lacked a discernible cardiac-frequency peak in *V*_downstream_ and was analyzed separately in Fig. S2 (see Methods).

We then defined percentile values for *V*_downstream_ for all ROIs: 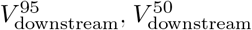, and 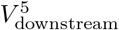 as the 95th, 50th (median), and 5th percentiles (Fig. 2d). Across all mice we found a mean 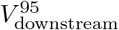 value of 121 µm/s for the straight PVS ROIs, demonstrating that the peak CSF velocity is much larger than the mean. Probability density functions of *V*_downstream_ for all 15 mice are included in Fig. S4. We obtained a full FOV PI of 3.9 ± 0.4 (Fig. 2e), which is considerably larger than the cardiovascular PI found in cerebral arteries (around 0.44) [45]. Higher PI are indicative of stronger particle oscillations and show a larger time varying response in the flow. We also investigated a PI based on the cross-stream component of velocity (Fig. S5) which is significantly lower than the downstream PI, is largest in straight PVS ROIs, and is weakly correlated with transverse motion of the artery across the cardiac cycle (transverse arterial motion is described below).

For different ROIs, straight PVSs had the highest mean PI at 6.2 ± 0.8, with some cases exceeding 10 (Fig. 2f). Lateral ROIs showed the smallest mean PI (2.5 ± 0.4), and the bifurcation PVS PI was 4.7 ± 1.2. PI was moderate for the full FOV ROIs (3.8 ± 0.4) due to spatial averaging. We checked that none of the reported PIs were erroneously inflated due to a small denominator in equation (3). Examining the statistical relationship between PI and mean flow speed showed negative trends for all ROIs. Bifurcation PVS ROIs showed the strongest negative association (*ρ* = − 0.943, Holm-adjusted *p* = 0.0667) suggesting that at slower flow speeds the pulsatile component of the flow is enhanced relative to the steady component (Fig. 2g).

These results show that high frame rate imaging resolves the transient CSF velocity peaks associated with each cardiac cycle, enabling quantification of a CSF PI akin to cardiovascular PI. While CSF PI is spatially heterogeneous and variable across mice, the pulsatility itself is strongly coupled to the cardiac cycle. Understanding the origin of this heterogeneity and whether it drives net transport requires considering both local arterial motion and the broader hydraulic properties of the CSF transport network.

#### Simulations indicate domain length is a critical parameter and net flow is not driven by cardiac pulsations

We idealized the PVS along the MCA as a single concentric circular annulus (Fig. 3a) and derived a reduced-order set of governing equations that conserve mass and momentum by applying the finite volume method (see Supplementary Materials §A). In this formulation, a prescribed arterial pulsation propagates along the inner boundary of the PVS with a speed of 1 m/s at a frequency of 5 Hz. We verified our simulation against an analytical solution [31] (Fig. S6a).

**Figure 3.**
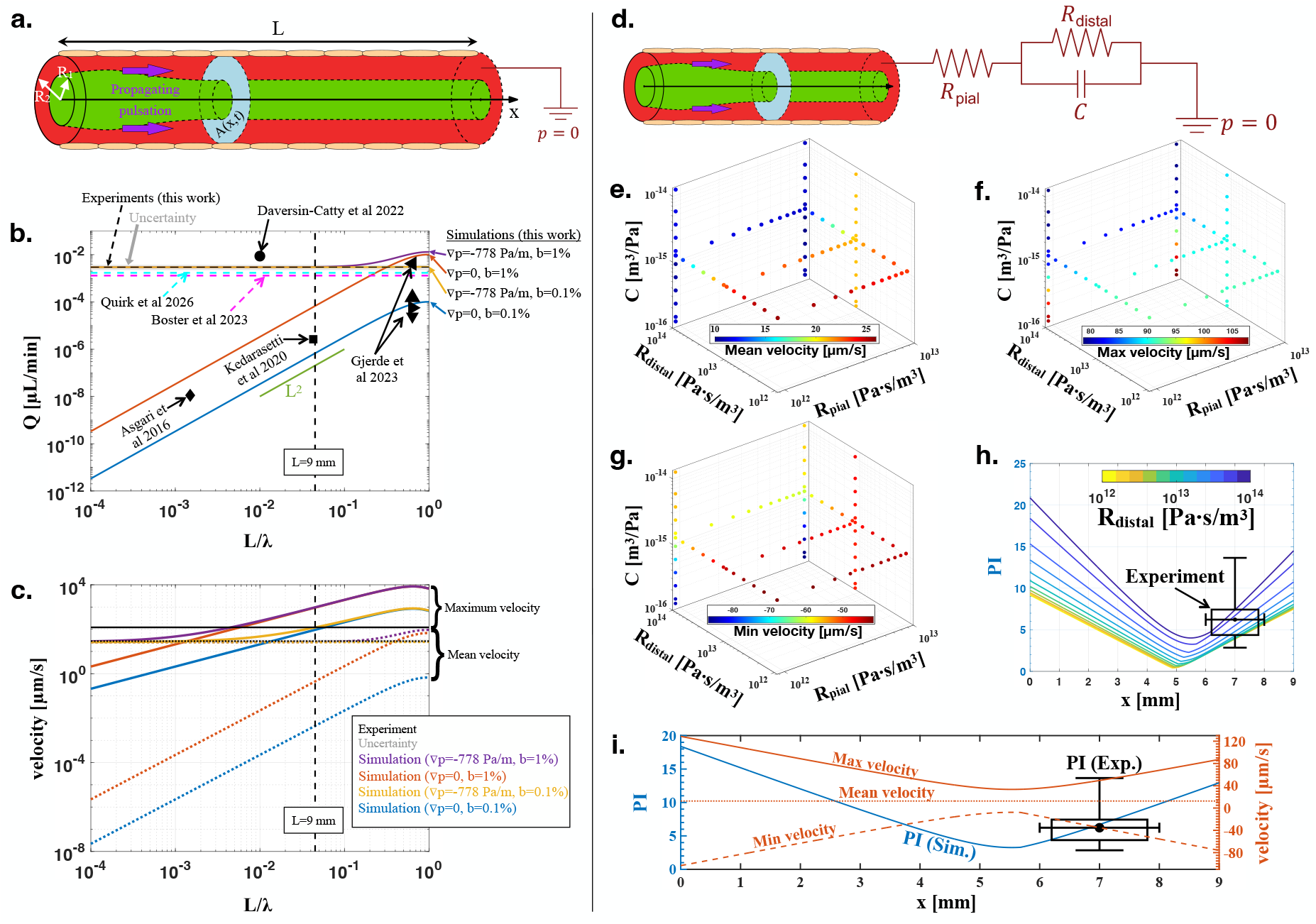
One-dimensional finite volume simulations indicate domain length is a critical parameter, and Windkessel boundary conditions alter flow speed and PI. **(a)** Schematic of CSF flow in a single PVS with variable domain length. **(b)** Comparison of time-averaged volume flow rate *Q* from simulations as a function of nondimensional domain length *L/λ*, compared with prior numerical studies [18, 19, 46, 32], prior experimental studies [5, 27, 40], and results reported here. **(c)** Plots of maximum and mean velocity for varying pulsation amplitude and pressure gradient. The minimum velocity is almost identical in magnitude (but negative) relative to the maximum velocity for zero pressure gradient (and it is symmetric about the mean velocity for a non-zero pressure gradient); we excluded it from the figure for the sake of clarity. **(d)** Schematic of the Windkessel boundary condition applied at the outlet of the PVS computational domain. **(e-g)** Variation of mean, maximum, and minimum CSF velocity (computed over space and time) with different Windkessel parameters. **(h)** PI as a function of domain location *x* for different values of *R*_distal_, which agrees well with our experimental measurements. *R*_pial_ = 8.8 × 10^12^ Pa ·s/m^3^ and *C* = 1.3 × 10^−15^ m^3^/Pa for all simulations. **(i)** Plot illustrating how the velocity (orange; right *y*-axis) varies along the domain in the simulation, giving rise to a spatially-variable PI (blue; left *y*-axis). *R*_distal_ = 6.3 × 10^13^ Pa·s/m^3^; *R*_pial_ and *C* are the same as Fig. 3h.

Simulations were conducted for sinusoidal wall motion amplitudes of *b* = 1% and *b* = 0.1%, and we examined the influence of domain length on the resulting volume flow rate *Q*. We found *Q* increases quadratically with *L* (green line in Fig. 3b), in agreement with analytical predictions [47, 46]. With zero external pressure gradient ∇*p*, the simulation results are consistent with previous computational studies of PVS flow driven solely by arterial wall motion [18, 19, 46]. Our results show the predicted *Q* varies substantially across previous studies largely because of differences in the simulated domain length. For example, Asgari et al. [18] simulated only a short PVS segment (approximately 150 µm), resulting in very small *Q*. In contrast, Kedarasetti et al. [19] considered a much longer domain (~ 5 mm), leading to substantially larger *Q*. Both values they report are in the range that we compute between *b* = 0.1% and *b* = 1%, but they fall three or more orders of magnitude below experimentally measured values (Fig. 3b). Values from Gjerde et al [46] are also included which simulate flow driven by vasomotion (not cardiac pulsation), which has a much shorter wavelength. We looked for evidence of vasomotion in our experiments but found inconsistent power across mice in the relevant range of frequencies.

We next tested how transport changes when an external pressure gradient ∇*p* is implemented. For ∇*p* = −778 Pa/m, the model reproduces the experimental *Q* measured in this study (Fig. 3b). When the domain size is much smaller than the wavelength (*L/λ* ≲ 0.1), the external pressure gradient dominates the net flow (purple/yellow versus orange/blue curves in Fig. 3b), consistent with Sharp et al. [48]. Notably, when *L/λ* ≳ 0.1 (for *b* = 1%), cardiac pulsations substantially increase *Q* (purple curve in Fig. 3b). However, we estimated that for mice the PVS length is about *L* = 9 mm [49] (see Methods), while the pulse wavelength is *λ* = *c/f* = (1 m*/*s)*/*(5 Hz) = 0.2 m, yielding *L/λ* = 0.045 (vertical dashed line in Fig. 3b). For the human case, assuming an M1 length of *L* ≈ 16.5 mm [50], a pulse wave speed of *c* ≈ 5 m*/*s, and a cardiac frequency of *f* ≈ 1 Hz gives *λ* ≈ 5 m and *L/λ* ≈ 0.0033. Therefore, the human MCA segment represents an even smaller fraction of the arterial pulse wavelength than in mice, and idealized simulations would similarly suggest the net flow is driven predominantly by an external pressure gradient in both species. A similar conclusion was reached by Daversin-Catty et al [32], also plotted in Fig. 3b. Notably, without an external pressure gradient, wave amplitude significantly affects *Q* (see separation between orange and blue curves in Fig. 3b). In contrast, when substantial ∇*p* is present, the *Q* sensitivity to pulsation amplitude is reduced (see smaller separation between purple and yellow curves in Fig. 3b). Fig. S6b provides a zoomed-in comparison between experiment and these simulation cases.

The nondimensional domain length also has a major impact on the velocity (Fig. 3c). The mean velocity (dotted lines) varies with a similar functional form to that of *Q* since they are linearly proportional. For the maximum velocity, the trend is similar to that of the mean velocity. However, the effect of the external pressure gradient on the maximum velocity is more pronounced at small *L/λ* and becomes negligible as *L/λ* increases. In particular, for fixed *b*, the maximum velocity curves merge at much lower values of *L/λ* than the mean velocity or *Q*. Unlike the mean velocity, which increases monotonically with *L/λ*, the maximum velocity reaches a peak at *L/λ* = 0.628 and then decreases slightly.

In summary, finite volume simulations with a realistic domain length predict: (i) net flow is generated by an external pressure gradient of at least −778 Pa/m, (ii) cardiac pulsations contribute negligibly to net flow, and (iii) cardiac pulsations determine the peak CSF velocity. Our simulations with a pulsation amplitude of *b* = 0.1% matched experimental measurements most closely (Fig. 3c). However, Mestre et al [5] showed that cardiac pulsation amplitude is around 1%. Hence, these idealized simulations do not accurately capture the dampening of peak CSF velocity, likely due to unrealistic boundary conditions, at least in part.

#### A realistic Windkessel boundary condition improves simulation agreement with experiment and helps explain experimental variation in PI

In the cardiovascular modeling literature, Windkessel boundary conditions are well-established and widely implemented as an effective approach for capturing the hydraulic resistance and compliance of downstream vasculature beyond the primary computational domain [51, 52], yet very few numerical simulations of CSF flow have implemented such boundary conditions [53, 54]. We implemented a three-element resistor-capacitor-resistor (RCR) Windkessel boundary condition at the outlet of the 1D model (Fig. 3d) to more accurately model *in vivo* conditions. This RCR boundary condition is essentially a time-varying pressure boundary condition that regulates the flow. It has two resistance parameters, *R*_pial_ and *R*_distal_, which augment the resistance of the pial PVSs and the distal flow pathways (e.g., penetrating PVSs). The capacitance parameter, *C*, captures the compliance of downstream flow pathways.

We varied the Windkessel parameters over physiologically relevant ranges. Distal resistance and capacitance were each varied over two orders of magnitude, from 8.8 × 10^11^ to 8.8 × 10^13^ Pa·s/m^3^ [54] and from 1.3 × 10^−16^ to 1.3 × 10^−14^ m^3^/Pa, respectively. The latter values are smaller than the subarachnoid CSF compliance reported by Ladrón-de-Guevara et al. [54], as our reduced order model represents an idealized single PVS pathway without bifurcations or coupling to the broader CSF system. *R*_pial_ was varied from 8.8 × 10^11^ to 8.8 × 10^12^ Pa·s/m^3^, similar to values estimated by Tithof et al. [55].

Variations in Windkessel parameters significantly affect CSF velocity (Fig. 3e-g). Mean velocity (about 10 to 30 µm/s) is most sensitive to *R*_pial_ and *R*_distal_. Maximum velocity *V*_max_ (about 80 to 110 µm/s) depends primarily on compliance, while minimum velocity *V*_min_ (about −90 to −40 µm/s) is influenced by all three Windkessel parameters. We used these values to compute a simulation PI:

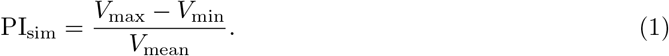

Here, mean velocity is used instead of median velocity in the denominator, as it is more robust and exhibits less fluctuation in the simulation. We find that PI_sim_ varies substantially over space and depends most strongly on *R*_distal_ (Fig. 3h); see Fig. S7a-b for dependence on other parameters. At any given *x*-location, PI_sim_ changes more than two-fold over the range of *R*_distal_ values tested (Fig. 3h). To make comparisons to the experiment, we estimated that our *in vivo* imaging was performed at *x*≈ 7 ± 1 mm along the MCA (see Methods); the spread in PI_sim_ due to different *R*_distal_ values thus agrees well with the range and uncertainty of experimental PI plotted in Fig. 3h. Simulations also predict that PI_sim_ reaches a minimum near the center of the domain (approximately 5–6 mm) and is higher toward both ends. This is because the domain is much shorter than the wavelength, so the oscillating PVS volume draws fluid in and expels fluid out at the two ends, producing the largest oscillatory flow amplitude there. Increasing *R*_distal_ elevates PI_sim_ throughout the domain, with the strongest effect near the inlet and outlet. In contrast, varying *R*_pial_ or *C* has a small effect on the local value of PI_sim_ (Fig. S7c-d).

Overall, these results suggest that variation in *R*_distal_ strongly modulates PI_sim_, leading to a range of values that agree well with experiments. In particular, larger values of *R*_distal_ agree better with the mean PI from experiments (Fig. 3i). We varied simulation parameters in an attempt to match experimental mean values; we obtained a mean velocity of 44.6 µm/s and a maximum velocity of 125 µm/s for an external pressure gradient of −2667 Pa/m, a pulsation amplitude of *b* = 0.22%, and Windkessel boundary conditions of *R*_distal_ = 6.3 × 10^13^ Pa·s/m^3^, *R*_pial_ = 8.8 × 10^12^ Pa·s/m^3^, and *C* = 1.3 × 10^−15^ m^3^/Pa.

### Heterogeneity of phase between arterial wall velocity, CSF velocity, and transverse wall motion

Mestre et al. [5] previously showed that CSF flow in pial PVSs pulses in synchrony with the cardiac cycle. That study collected 2D images using 2PM to measure CSF flow, followed by separate 2PM line scans (at about 1 kHz) to measure vessel diameter. Here, we measured vessel diameter by directly analyzing 2D images from 2PM, which has the benefit that vessel diameter and CSF velocity can be measured simultaneously and directly compared (Fig. 4a). We also tracked the transverse motion of the artery, which a prior numerical study [32] suggested can impact CSF flow, rendering the dynamics complex and nuanced. The signals we obtained are noisy, especially vessel diameter (due to the much lower imaging frequency compared to line scans), but all power spectra for all signals for all 15 mice possess a clear peak at the cardiac frequency, except for that of *V*_downstream_ for one mouse (Fig. S2e,f and Fig. S3c,f) and the transverse motion signal for one other mouse. We computed and plotted the dominant Fourier mode for each signal (thick lines in the top three panels of Fig. 4b).

**Figure 4.**
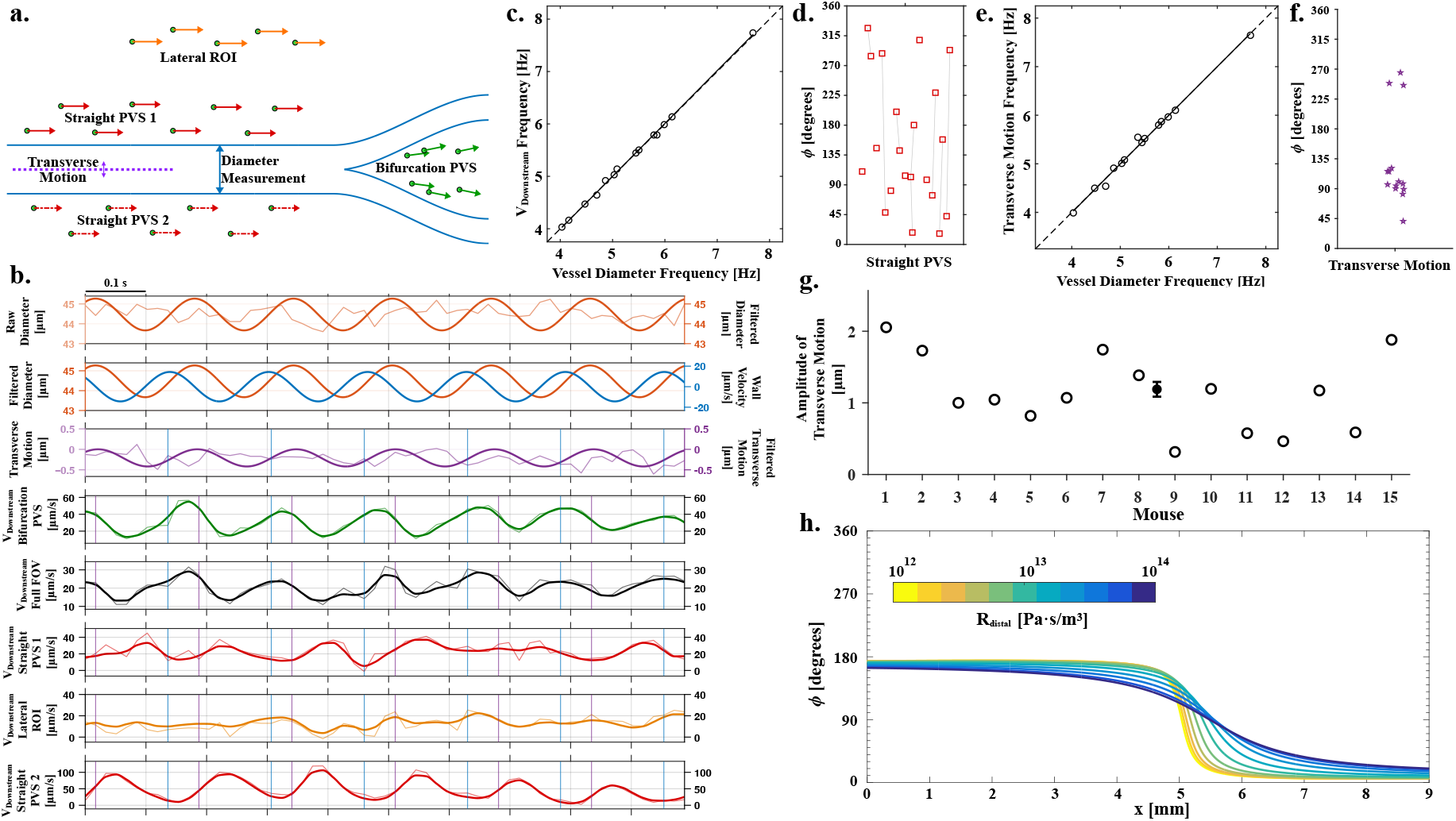
Phase analysis demonstrates that CSF velocities are synchronized with arterial wall velocity through both radial (dilation/constriction) and transverse arterial motion while also being spatially heterogeneous. **(a)** Schematic illustrating the analyzed PVS ROIs, arterial diameter changes due to cardiac pulsations, and arterial transverse motion all used for subsequent phase analysis. **(b)** Raw (thin curves) and Fourier filtered (thick curves for top three panels) or smoothed (thick curves for *V*_downstream_ panels) time-synchronized signals including arterial diameter, arterial transverse motion, and *V*_downstream_ for different ROIs from mouse 3. **(c)** Plot of the dominant *V*_downstream_ frequency versus the dominant frequency from vessel diameter measurements; the dashed line indicates *y* = *x* and solid line indicates the linear regression (*R*^2^=0.99). **(d)** Phase offset *ϕ* between the dominant frequencies of the arterial wall velocity and *V*_downstream_ in Straight PVS ROIs (where each phase offset value quantifies how much the peak in *V*_downstream_ lags the arterial wall velocity peak in time) for 21 Straight PVSs across *n* = 13 mice. Light gray lines indicate Straight PVS ROIs from the same mouse. **(e)** Plot of the dominant transverse motion frequency versus the dominant frequency from vessel diameter measurements; the dashed line indicates *y* = *x* and solid line indicates the linear regression (*R*^2^=0.99). **(f)** Phase offset between the dominant frequencies of the arterial wall velocity and transverse motion for *n* = 15 mice. **(g)** Amplitude of arterial motion in the transverse direction; the error bar indicates mean ± SEM, *n* = 15 mice. **(h)** Plot of the phase between the arterial wall velocity and the peak CSF velocity from the 1D finite volume simulations, computed over space and for different values of *R*_distal_. *R*_pial_ = 8.8 × 10^12^ Pa·s/m^3^ and *C* = 1.3 × 10^−15^ m^3^/Pa for all simulations.

From Fourier analysis, we confirmed that the dominant arterial pulsation frequency (5.34 ± 0.25 Hz on average) closely matched that of *V*_downstream_ (5.34 ± 0.26 Hz on average) and was in the expected heart rate range for mice under KX anesthesia [56] (Fig. 4c). These measured frequencies were very strongly correlated, with the linear regression yielding a coefficient of determination of *R*^2^ = 0.99. From the dominant arterial pulsation frequency, we analytically computed the associated arterial wall velocity (blue curve in Fig. 4b), which we then used to compute its relative phase *ϕ* with the different *V*_downstream_ ROIs (see Methods). We found *ϕ* varied substantially across different mice and even different Straight PVSs within the same mouse (Fig. 4d). We were surprised by this finding, so we investigated the variation of *ϕ* across different segments of each time series and found good temporal consistency in phase, with a standard deviation of 35^◦^ or less across all included mice.

We also analyzed the transverse motion of the artery (purple curve in Fig. 4b), which yielded a mean amplitude of 1.06 ± 0.14 µm (Fig. 4g). We performed Fourier analysis again and found that the dominant frequency of the arterial transverse motion closely matched that of the vessel diameter, coinciding with the cardiac frequency range (Fig. 4e). The phase *ϕ* between the arterial wall velocity and transverse motion clearly exhibited a bimodal distribution localized near 90^◦^ and 270^◦^ (Fig. 4f). The majority of mice exhibited anterior transverse motion lagging a quarter cycle (~ 90^◦^) after the arterial wall velocity. Since the phase between arterial diameter and wall velocity is 90^◦^, transverse motion is therefore approximately in phase or 180^◦^ out of phase with arterial pulsations, i.e., systole and diastole coincide with the maximum arterial transverse displacement in one direction or the other.

We also performed 1D finite volume simulations with Windkessel boundary conditions to investigate the phase relationship between *V*_downstream_ and arterial wall velocity (the 1D simulations cannot capture transverse motion of the artery). Simulations demonstrate that the relative phase between *V*_downstream_ and arterial wall velocity varies substantially but monotonically along *x*; furthermore, the value of the phase is fairly sensitive to *R*_distal_ (Fig. 4h). The phase also varies with pulsation frequency, *R*_pial_, and *C* (Fig. S7e-h), but much less so than with *R*_distal_. However, the simulations do not capture the extent of spread in phase observed in experiments.

In summary, the phase between arterial wall velocity and CSF flow in Straight PVSs measured experimentally is variable across mice and even for different PVSs within the same mouse. Simulations suggest some modest differences in phase may be explained by variable imaging location and down-stream hydraulic properties, especially *R*_distal_, but simulations do not predict the phase to ever be above 180^◦^. Further analysis is needed to evaluate whether transverse arterial motion may contribute to variable phase between arterial wall velocity and *V*_downstream_.

### In vivo measurements and computational simulations characterize spatiotemporal variation in glymphatic wall shear stress

Mechanotransduction refers to the process by which cells convert mechanical forces, such as WSS, into biochemical signaling. Recent studies have shown that Piezo1, which is expressed in both endothelial cells and astrocyte endfeet, is linked to neurogenesis and cognitive function [57], blood-brain barrier disruption following ischemic stroke [58], and even amyloid-β clearance via enhanced phagocytosis and lysosomal activity [59]. To estimate mean and peak WSS in space and time along the PVS boundaries, we performed mouse-specific finite element simulations of fluid flow through an idealized cross-sectional PVS geometry (Fig. 4a), parameterized by our arterial diameter and PVS width measurements. We performed dimensionless finite element simulations, we scaled the results by the experimentally measured velocity (either the temporal mean or maximum [i.e., 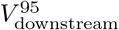]), then computed and plotted the WSS (Fig. 5b; see Methods). The WSS varies in space for both boundaries, with maximum values occurring at different angular locations (Fig. 5c). The temporal mean and maximum WSS, owing to the pulsatile nature of the flow, is plotted in Fig. 5d-e. We computed a temporal maximum WSS of 0.064 Pa on the arterial wall (0.027 Pa averaged over the arterial wall) and 0.050 Pa on the outer PVS boundary (0.030 Pa averaged over the outer wall).

**Figure 5.**
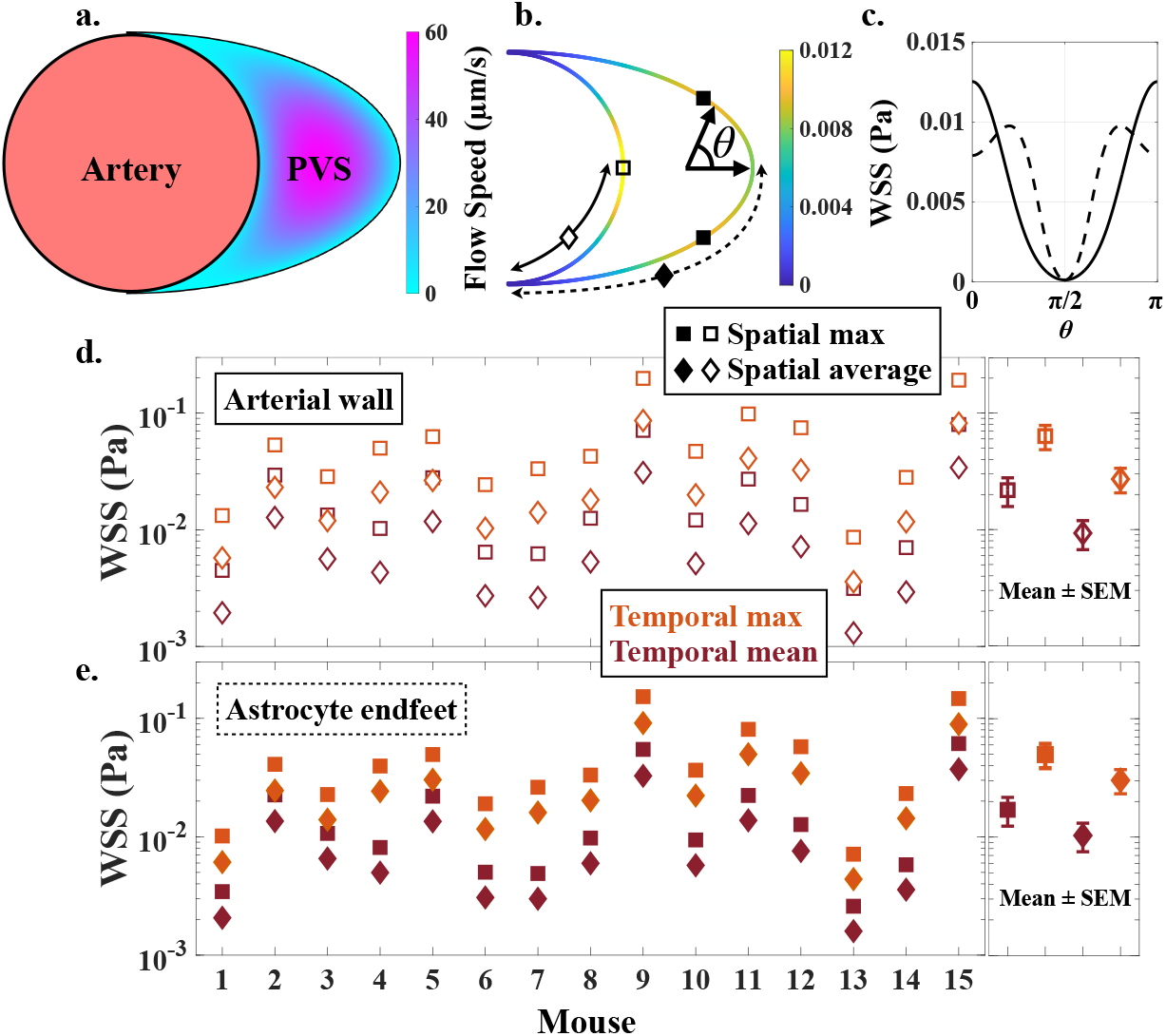
Mouse-specific finite element simulations provide estimates of mean and maximum wall shear stress (WSS) in time and space. **(a)** Flow speed profile in an idealized circle-ellipse model of a PVS [36], and **(b-c)** the associated WSS for mouse 8. **(d-e)** WSS on the arterial wall (open symbols) or on the astrocyte endfeet forming the outer PVS boundary (solid symbols). Squares correspond to the maximum value in space, diamonds correspond to the spatial average, orange is scaled by the temporal maximum speed, and burgundy is scaled by the temporal mean speed. Error bars indicate mean ± SEM, *n* = 15 mice.

## Discussion

Numerous studies demonstrate that reductions in glymphatic flow coincide with development of neurodegenerative pathology in both mouse models [60, 61, 62, 63] and patients [64, 65, 66]. While the direction of causality has not been fully established, recent studies have shown that acutely reducing glymphatic flow leads to impaired solute clearance from the brain [67, 68, 69]. *In vivo* measurements reported in this study provide the most thorough characterization of CSF flow in healthy, young mice, detailing the high level of flow heterogeneity, which provides nuanced and important details for future studies seeking to characterize changes in glymphatic CSF flow for a variety of neurological diseases. In particular, we found that flow through Straight PVS regions is approximately three times faster than flow in bifurcation regions (Fig. 1c); Schain et al [37] noted that bifurcation regions have larger PVS cross-sectional areas, which may explain the slower flow in those regions due to mass conservation (i.e., fixed *Q* = *V A* along a PVS implies that a larger cross sectional area *A* in some region leads to smaller local velocity *V*). Notably, we showed that flow exists in some lateral regions with an average speed of 25. ± 1 3.3 µm/s demonstrating that appreciable advective transport can occur away from the primary PVS, at least at the surface of the brain. However, this should not be interpreted as interstitial flow through the brain parenchyma. Rather, these flows are possibly located in the narrow space between the parenchyma and the membrane forming the upper boundary of the pial PVS (e.g., see Fig. 2f in or Fig. 2c in [70]). The flows in this region are directional and exhibit significantly reduced PI (Fig. 2g), indicating net pressure gradients are appreciable and adjacent compliant tissues dampen the pulsatile component of the flow.

We achieved a 2PM imaging frame rate (113 Hz) higher than any prior CSF studies allowing us to resolve the pulsatile velocity peaks in Straight PVSs (121 ± 16.7 µm/s) associated with each cardiac cycle better than prior studies performed at 30 Hz which reported peaks around 30 to 80 µm/s for individual mice [5, 26, 27]. Resolving these peak velocities was essential for reliably estimating the CSF PI and peak WSS. Quantifying these two variables, as well as the phase offset between arterial wall velocity and CSF flow, has important implications for both the design of future studies and the interpretation of prior publications, as discussed below.

Our results suggest CSF PI provides novel information about glymphatic function beyond just mean flow speed. While mean flow speeds or volume flow rates reflect how much CSF is transported through the network, PI characterizees the hydraulic network’s response to arterial pulsations and is influenced by distal resistance and compliance. Because CSF cannot currently be measured deep within the brain with high spatial and temporal resolution, surface measurements of PI may provide a sensitive marker for distal hydrodynamic alterations that lead to glymphatic impairments. Future studies could leverage the optical imaging approach demonstrated here to infer changes in glymphatic compliance and/or resistance associated with neurodegeneration, traumatic brain injury, brain edema, changes in aquaporin-4 polarization/expression, PVS dilation, or many other age-related pathologies.

An additional advantage of PI is that it characterizes a component of CSF dynamics that may be biologically important but distinct from net transport. Our simulations suggest cardiac pulsations determine peak CSF velocities, whereas the mechanisms governing net CSF flow may differ. Because peak velocity determines peak WSS on the PVS boundaries, altered cardiac pulsatility could affect mechanotransductive signaling even without substantial change in net flow. Cibelli et al [71] reported astrocytic calcium signaling for WSS thresholds below 0.01 Pa, comparable to our spatially and temporally averaged WSS of 0.010 Pa on the brain-facing PVS boundary. Prior studies suggest this boundary is formed by astrocytic glia limitans superficialis and its associated basement membrane [72, 73]. Wakida et al [74] found that astrocytic phagocytosis is enhanced at a WSS of 0.025 Pa, slightly below our measured spatially-averaged temporal peak in WSS (0.030 Pa). Our computed temporal and spatial maximum of 0.050 Pa is considerably higher; while prior work has shown that transient WSS elevation has a weaker signaling effect than constant WSS elevation [75], it is unclear to what extent mechanotransduction will occur for a periodic stimulus generated by cardiac pulsations.

Our measurements of the phase *ϕ* between arterial wall velocity and downstream CSF velocity revealed substantial variability across mice and PVSs, despite consistency of *ϕ* over the signal’s time series. This disagrees with Mestre et al. [5] and has important implications for testing the feasibility of various CSF driving mechanisms proposed in prior numerical studies. Mestre et al. [5] showed that the temporal variation in root-mean-square velocity Δ*v*_*rms*_ peaks in approximate synchrony with the arterial wall velocity (their Fig. 3f). However, this measurement is based on a non-negative quantity that includes the cross-stream component of the velocity (which will generally be maximum when the wall velocity is maximum). Analysis of *V*_downstream_, presented here, is a more proper approach as it considers the net flow component which is used in numerical investigations of phase. Peristalsis models (without Windkessel boundary conditions) [76, 14, 19] predict *ϕ* = 270^◦^, which reasonably agrees with measurements for a subset of our mice, but these studies assume a non-physical domain length *L* = *λ* and no downstream impedance. Models combining PVS-width changes with transmantle pressure waves predict phases of 108^◦^ and about 180^◦^, respectively [20, 30]. Our measurements do not strongly support or refute this hypothesis. Two other recent studies have proposed that astrocyte endfeet may function as valves, generating directional flow from oscillatory pressure in the PVS [77, 78]. Gan et al [78] derived an analytical expression for the phase, which predicts a lag that varies between 0^◦^ and 90^◦^, potentially aligning with about half of our measurements. Kedarasetti et al [19] showed that using a realistic PVS shape, domain length, and arterial pulsation waveform led to a phase of approximately 330^◦^ (estimated in [54]). Ladrón-de-Guevara et al [54] applied a two-element Windkessel boundary condition to a model of PVS flow, and they concluded that doing so shifts the phase in these prior studies such that the CSF flow is almost synchronous with wall velocity (*ϕ* ≈ 0^◦^, as described by Mestre et al [5]). Our experimental measurements generally disagree with predictions from these two studies. Notably, Ladrón-de-Guevara et al [54] implemented a two – not three – element Windkessel boundary condition and did not consider variation of phase with axial location or Windkessel parameters. Finally, it is unclear whether arterial transverse motion may influence the phase between arterial wall velocity and *V*_downstream_; in either case, transverse artery motion likely contributes negligibly to net transport, but it may increase mixing in PVSs [79]. Further numerical investigations – which account for spatial variation in phase and dependence on Windkessel boundary conditions – are needed to further constrain the range of potential driving mechanisms.

This study has several limitations. All experiments were performed under KX anesthesia, not natural sleep. Emerging evidence [23] suggests glymphatic transport is greatest during sleep due to vasomotion, characterized by large, low fequency amplitude fluctuations in arterial diameter that modles suggest may drive appreciable net flow [46]. Importantly, power spectra from our arterial diameter and CSF velocity measurements showed inconsistent power in the range of frequencies associated with vasomotion. Quantitative measurements of CSF flow and PI under wake and sleep conditions would provide a valuable framework for testing whether PI is predictive of changes in downstream resistance (the expansion of the extracellular space during sleep [80] is estimated to cause a 5.5-fold decrease in hydraulic resistance in the brain parenchyma [55]). An additional limitation is that our finite volume simulations model the glymphatic pathway as one single PVS, not a network of branching PVSs. We cannot dismiss the possibility that arterial pulsations propagating across a realistic, branching network may generate results quite different from those presented here. However, our modeling approach neglected both PVS branching and axial attenuation in PVS size which have opposing effects on net CSF flow which likely increase the robustness of our modeling. Specifically, branching tends to increase the total PVS cross-sectional area while attenuation decreases it.

## Methods

### Animals and surgical preparation

All experiments were approved by the University of Minnesota’s Institute for Animal Care and Use Committee (Protocol No. 2209-40409A) and were designed in consultation with the committee. 15 male C57BL/6 mice, 8-28 weeks old (The Jackson Laboratory), were anesthetized with ketamine-xylazine (100/10 mg/kg, intraperitoneally) along with Meloxicam (10 mg/kg, subcutaneously). Throughout surgery and imaging, the body temperature was maintained at 37.2 ^◦^C with a rectal probe-controlled heated platform (Physilab TCAT-2LV Controller). Removal of the skin and periosteum was performed to expose the skull, and optical access to the brain was created by implanting a cranial window over the MCA vascular territory on the right anterolateral parietal bone. Great care was taken to thin the skull path with a dental drill by drilling short paths in small layers for brief periods of time and wetting the skull area with saline between passes to not heat up the brain tissue. This process was done until the drilled path was approximately 10 µm, allowing safe excision of the skull piece while ensuring that the underlying dura mater was left intact. The cranial window was sealed with agarose (0.8% at 37 ^◦^C) underneath, and dental cement was applied along the coverslip circumference to prevent loss of intracranial pressure. 2,000 kDa FITC dextran was introduced into the bloodstream through a retro-orbital injection [81]. A custom cannula was inserted into the cisterna magna which facilitating delivery of red fluorescent polystyrene microspheres (Invitrogen FluoSpheres 1.0 µm, 580/605 nm) to the CSF. The microspheres were briefly sonicated prior to infusion and were then injected into the cisterna magna at 2 µl/min for 5 min using a syringe pump.

### In vivo two-photon laser scanning microscopy

Two-photon imaging was performed on a Nikon A1RHD MP using a Plan Apo LWD 25x water-immersion objective (NA 1.1) and resonant scanners. A Mai Tai eHP tunable IR laser was tuned to 870 nm to excite both the 2,000 kDa FITC dextran in the blood and the red fluorescent microspheres in the CSF. Emitted fluorescence from FITC dextran and red microspheres was detected using non-descanned GaAsP detectors through 510/80 nm and 607/70 nm emission filters, respectively. Images were acquired at a spatial resolution of 1.02 µm/pixel and variable temporal resolution depending on the spatial dimensions. Conventional 2PM systems commonly use a resonant scanner for rapid scanning along one axis and a galvanometric scanner for the orthogonal axis. Thus, by reducing the field of view (FOV) to one-half (512 × 256 pixels) or one-quarter (512 × 128 pixels) along the slow (galvanometric) scanning direction (Fig. 1e-g), the frame rate can be approximately doubled or quadrupled from the standard 30 Hz used in prior studies [5, 26, 27]. This allows for much higher temporal resolution enabling higher fidelity particle velocity measurements in time that more accurately resolve the flow pulsatility. We tested frame rates as high as 720 Hz, but doing so necessitated a very narrow FOV in which anatomy was difficult to distinguish; additionally, such a narrow FOV led to sparse velocity measurements if the PVS was not perfectly aligned with the extended direction. In our presented work, we collected our datasets at: 30 Hz (1 mouse), 58 Hz (5 mice), 113 Hz (4 mice), both 58 Hz and 113 Hz (4 mice), and both 113 and 226 Hz (1 mouse). Imaging was performed along the MCA and its associated branches, which were identified by locating the anterolateral aspect of the cranial window and following the vessel toward the midline. The focal plane was adjusted in the axial imaging direction to maximize the apparent diameter of the primary arterial segment within the field of view, thereby positioning the imaging plane as close as possible to the vessel centerline.

### Particle tracking velocimetry

Fluorescent microspheres flowing in pial PVSs were tracked using a robust and well-established custom particle tracking velocimetry (PTV) Matlab code [5, 26, 35], with one important improvement. We extended the code to track particles based on fitting the intensity profile to a 2D Gaussian profile rather than using Matlab’s “regionprops” function (the latter computes the centroid based on the pixel arithmetic mean). We generated and analyzed synthetic images (not shown) to verify that this change led to reliable improvements in tracking. Lagrangian tracks from PTV store particle positions and velocities throughout time. We created a Cartesian grid over the entire FOV for each recording, which we used to bin then compute average (Eulerian) velocity fields and speed maps (Fig. 1e-f). Stagnant particles were masked by computing dynamic background images [35] to prevent underestimating CSF flow. User-defined ROIs were drawn then used to mask speed maps to compute mean flow speeds in different regions (Straight PVSs, Lateral ROIs, etc.). Higher imaging frame rates were achieved by reducing the FOV along the galvanometric scanning direction (described above). Although increasing the imaging frame rate can modestly increase localization-related uncertainty in PTV, these errors are generally small compared with the benefits of improved temporal resolution for rapidly accelerating particles [82].

### Downstream velocity and pulsatility index

We quantified the instantaneous component of CSF velocity in the downstream direction as:

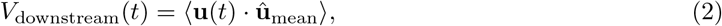

where **u**(*t*) is the instantaneous velocity of a particle at time *t*, ***û***_mean_ is the nearest unit vector of the net transport determined from the time-averaged velocity field, and ⟨·⟩ denotes averaging over all particles tracked at time *t* for a given ROI.

Also within that ROI, the pulsatility index (PI) was computed as:

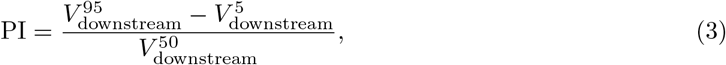

where the superscript *x* denotes the *x*^th^ percentile. Percentiles were used to avoid outliers that might occur due to erroneous tracks in the experimental measurements.

### Power spectrum analysis

Downstream velocity (from PTV) and both arterial diameter and transverse motion (from custom Matlab vessel tracking codes) were processed using Matlab’s “fft” function. The associated power was computed and plotted either as a function of frequency (for a single 60 s time series) or as a spectrogram (for several 60 s time windows shifted across the entire recording). One dataset (mouse 9) exhibited an unusually high mean flow speed and lacked a discernible cardiac-frequency peak in the *V*_downstream_ power spectrum. This dataset was therefore excluded from analyses in Figs. 1-2 and was analyzed separately in Fig. S2.

### Phase analysis

Phase and coherence between arterial wall velocity and CSF downstream velocity (*V*_downstream_) or transverse arterial motion were quantified using cross-spectral analysis at the cardiac frequency. Signals were divided into overlapping Hann-windowed segments with 50% overlap, and only segments containing continuous data were included. A default window duration of 10 s was used. Each segment was detrended and Fourier transformed, and the cardiac frequency was identified from the arterial diameter spectrum. Arterial wall velocity was obtained from the diameter signal in the frequency domain as *W* (*f*) = *i*2*πf D*(*f*). The wall-velocity autospectrum (*S*_*ww*_), comparison-signal autospectrum (*S*_*vv*_), and cross-spectrum (*S*_*wv*_ = *W* ^∗^*V*) were averaged across all valid segments. Magnitude-squared coherence was calculated as:

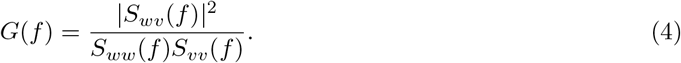

Coherence, which ranges from 0 to 1, was used to quantify the consistency of the phase relationship between the signals at the cardiac frequency. Phase estimates with *G <* 0.3 were excluded from subsequent phase summaries to limit inclusion of poorly coupled signals. Phase was calculated as *ϕ*(*f*) = arg[*S*_*wv*_(*f*)] and expressed in the range [0^◦^, 360^◦^), such that *ϕ* represents the phase of the CSF velocity or transverse-motion signal lagging arterial wall velocity. For consistency, we defined the positive transverse direction of the artery based on anatomy so that it corresponded to the anterior direction (i.e., top of the 2PM image).

### PVS/artery area ratio

Artery diameter measurements (twice the radius *R*_1_) were determined for each dataset using a custom Matlab code that tracks the vessel edge via intensity thresholding of the FITC dextran fluorescent dye flowing in the bloodstream [35]. Vessel edges were tracked in time at the full imaging frame rate (typically 60 or 113 Hz). PVS width was obtained by superimposing all particle tracks obtained for a given recording then measuring the width of that region using a custom Matlab code (Fig. S1b). The PVS cross-sectional area was estimated by assuming the PVS is shaped like an ellipse surrounding a circular artery [36] and was thus calculated as 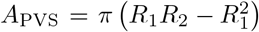, where *R*_2_ is the semi-major axis length, equal to *R*_1_ plus the PVS width. Experimental time-averaged volume flow rates *Q* were estimated from the mean flow speeds and *A*_PVS_ for each straight PVS. Importantly, our reported volumetric flow rate is through only one lobe of a pial PVS, as the straight PVS measurements (mean speed and PVS width) were always computed for one side of the MCA (Fig. S1b).

### Statistical analysis

All statistical analyses were performed in Matlab (version 24.2.0; The MathWorks Inc., Natick, MA). Data are presented as mean ± SEM unless otherwise indicated. For descriptive statistics, repeated measurements of the same ROI category within an individual mouse were averaged such that each mouse contributed a single summary value per ROI category. Comparisons among ROI categories were performed using linear mixed-effects models, with ROI and imaging frame rate included as fixed effects and mouse and recording included as random intercepts to account for repeated measurements. Normality of model residuals was assessed using the Lilliefors test. Statistical significance was assessed for fixed effects and pairwise ROI comparisons. Pairwise p-values were adjusted using the Holm method across all six possible ROI comparisons for each outcome. Frame rate was modeled as a continuous covariate in the primary analyses, with categorical frame-rate groups evaluated as a sensitivity analysis; the statistical conclusions were unchanged. Mouse-level paired Student’s t-tests or Wilcoxon signed-rank tests, as appropriate, were additionally performed as sensitivity analyses for comparisons between ROI categories present within the same mice. Associations between mean flow speed and PI were assessed using Spearman rank correlations, with Holm correction applied across the four ROI-specific correlations. Statistical significance was defined as Holm-adjusted *p <* 0.05 where multiple comparisons were performed.

### MCA length estimate

We estimated the murine MCA to be approximately 9 mm based on a digital graphical measurement we performed using a snapshot taken from the iDISCO brain vasculature reconstruction provided by Kirst et al [49] in their Video S1. We also used a snapshot from a dorsal view provided in this video to estimate (based on the coordinates of our cranial window implantation) that our measurement location was 2 ± 1 mm from the most distal pial branches of the MCA (i.e., 7 ± 1 mm from the base of the MCA).

### Numerical modeling of CSF flow in the PVS

Finite volume simulations: We idealized the PVS as the annular region between two concentric cylinders of radius *r*_1_ = 20 µm and *r*_2_ = 40 µm, where the inner wall undergoes time and space dependent expansion and contraction to represent arterial pulsations and the outer wall is assumed rigid. The flow is assumed to be incompressible and laminar, with viscosity *µ* = 1.0 × 10^−3^ Pa·s, and the numerical scheme was derived based on the finite volume method (see Supplementary Materials §A). Simulations were verified by comparing resutls to a 1D analytical solution [31] with a domain length *L* equal to one wavelength *λ* (Fig. S6a). Subsequent simulations imposed an external pressure gradient of either 0 or 778 Pa/m by specifying the inlet pressure as either 0 or − 778*L* Pa, respectively. The outlet boundary condition was either zero-pressure or a standard three-element Windkessel boundary condition with *R*_pial_ connected in series to *R*_distal_ and *C* connected in parallel.

Finite element simulations: Flow through straight PVSs was modeled by idealizing the PVS geometry as an annular domain with a circular inner boundary representing the artery and an elliptical outer boundary representing the surrounding tissue [36]. We assumed the flow is purely axial, and that it is laminar and fully-developed due to the small Reynolds and Womersley numbers [83]. Hence, the Navier-Stokes equation reduces to Poisson’s equation:

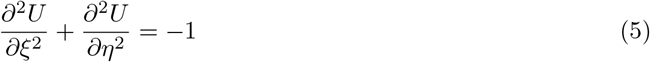

with dimensionless quantities 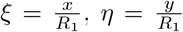, and 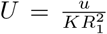, where *R*_1_ is the radius of the inner circular boundary and 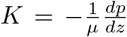 [36]. Equation (5) was solved using Matlab’s PDEToolbox. To dimensionalize results, dimensional volume flow rates *Q*_dim_ from experiments and the known fluid viscosity *µ* were used to compute the pressure gradient (or equivalently, the nondimensionalization constant *K*):

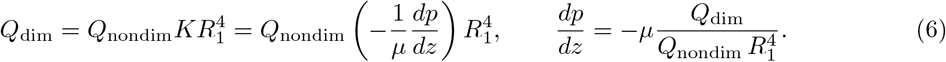

The corresponding pressure gradient values are reported in the Supplementary Material. The nondimensional wall shear stress was computed as ∂*U/*∂*n* at the boundaries of the PVS, where *n* is outward normal direction. The dimensional wall shear stress was computed as

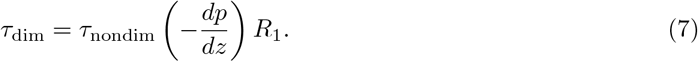

## Data availability

All two-photon images analyzed in this study are available via Zenodo at the following links:

https://doi.org/10.5281/zenodo.21984496

https://doi.org/10.5281/zenodo.21986687

https://doi.org/10.5281/zenodo.21987053

https://doi.org/10.5281/zenodo.21996238

https://doi.org/10.5281/zenodo.21999587

https://doi.org/10.5281/zenodo.22001103

https://doi.org/10.5281/zenodo.22003334

Measurements and numerical results presented in figures throughout this manuscript and simulation codes are included as Supplemental Data.

## Acknowledgments

This study was supported by the National Institute of Biomedical Imaging and Bioengineering and the National Center For Complementary & Integrative Health of the National Institutes of Health under Award Numbers R21EB036217 and R21AT013325, respectively. The content is solely the responsibility of the authors and does not necessarily represent the official views of the National Institutes of Health. This work was also supported by the Minnesota Office of Higher Education under Award No. 257252/3000008651.

Additional support was provided by the resources and staff of the University of Minnesota University Imaging Centers (UIC; RRID: SCR 020997). We thank Patrick Wiley, Jason Mitchell, Steve Schnell, Mary Brown, and Laurel Schuck for their technical assistance and guidance with two-photon imaging experiments. We also thank the Nedergaard and Hablitz Laboratories at the University of Rochester Medical Center and the Kodandaramaiah Laboratory at the University of Minnesota for their technical guidance and scientific expertise. The authors are particularly grateful to Michael Giannetto, Antonio Ladrón-de-Guevara, and Skyler Fausner for their mentorship and support.

## Author contributions

J.T., C.G., and K.H. conceived of the project. C.G. performed all mouse surgeries and imaging, with assistance from D.H.T., K.Di., S.S., T.R., K.Du., A.V., T.S.A. and D.K. All experimental data analysis was performed by C.G., J.T., D.H.T., K.Di., S.S., T.R., K.Du., A.V., and D.K. All statistical analysis was performed by C.G. All simulations and associated data analyses were performed by K.H. M.M. wrote the original version of the finite volume simulation codes. C.G., J.T., and K.H. wrote the manuscript. All authors approved of the submitted version.

## Supplementary Materials

**Figure S1.**
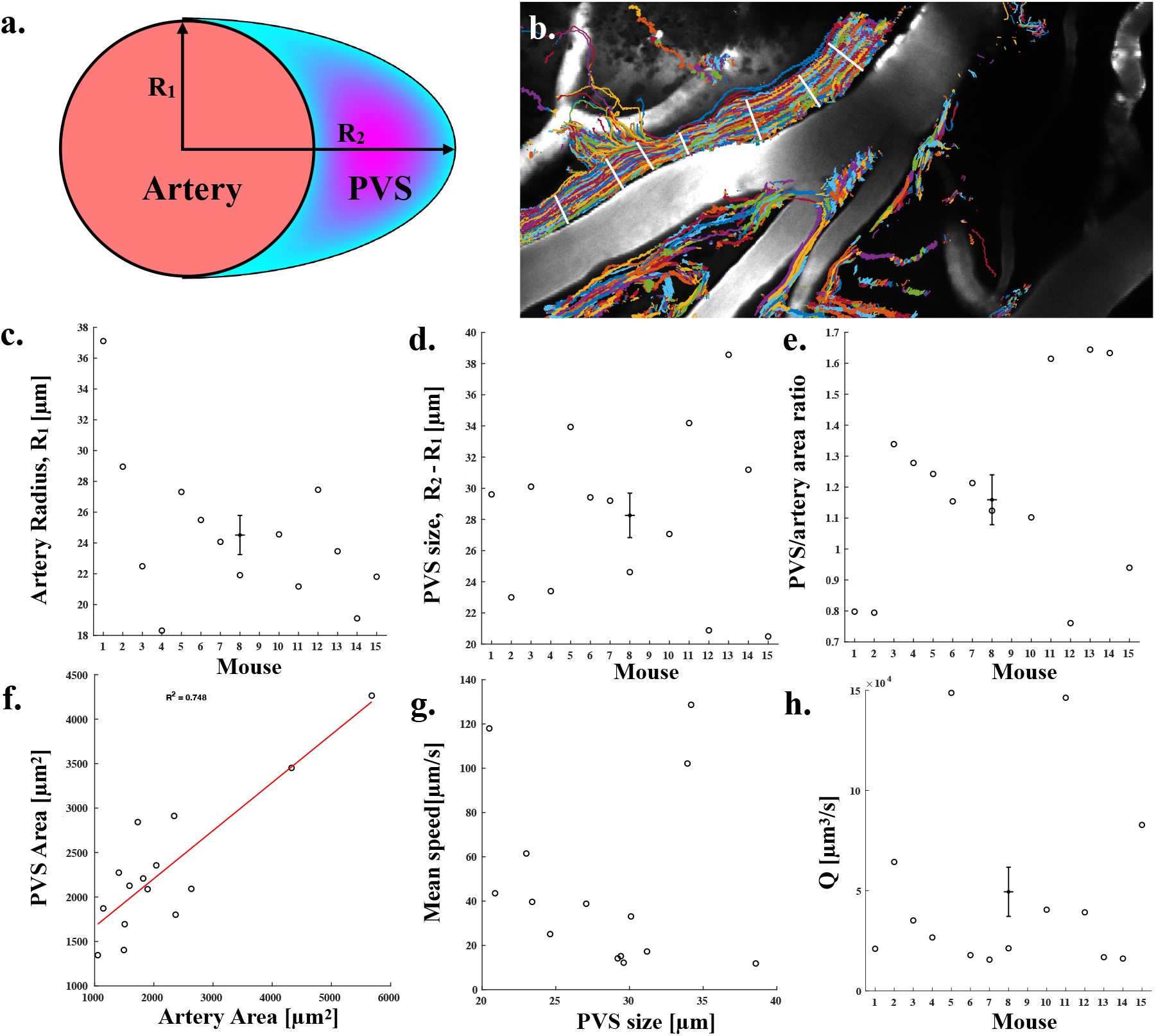
Experimental volumetric flow rates were estimated from measured mean speeds, and PVS cross-sectional areas were estimated from particle track widths and arterial diameter measurements. **(a)** Circle-ellipse idealized geometry [36] was used for estimating experimental PVS cross-sectional area and volumetric flow rates. **(b)** PTV tracks from each dataset were superimposed enabling estimates of *R*_2_ − *R*_1_ by measuring the width of the tracks. **(c)** Mean arterial radius for each mouse. The error bar near the center indicates the mean ± SEM. **(d)** Estimates of PVS size, *R*_2_ − *R*_1_, for each mouse. **(e)** PVS/artery area ratio for each mouse based on results shown in (c,d). This calculation assumes a two-lobe PVS area where *R*_2_ is the same for both PVS lobes. **(f)** PVS area linearly increases with arterial area with *R*^2^ = 0.75. **(g)** Mean speed in the PVS generally decreases with increasing PVS size with the exception of two outliers. **(h)** Experimental volume flow rates through a single PVS lobe on one side of the artery, estimated for each mouse. This was computed as the straight PVS mean speed times 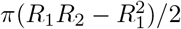.

**Figure S2.**
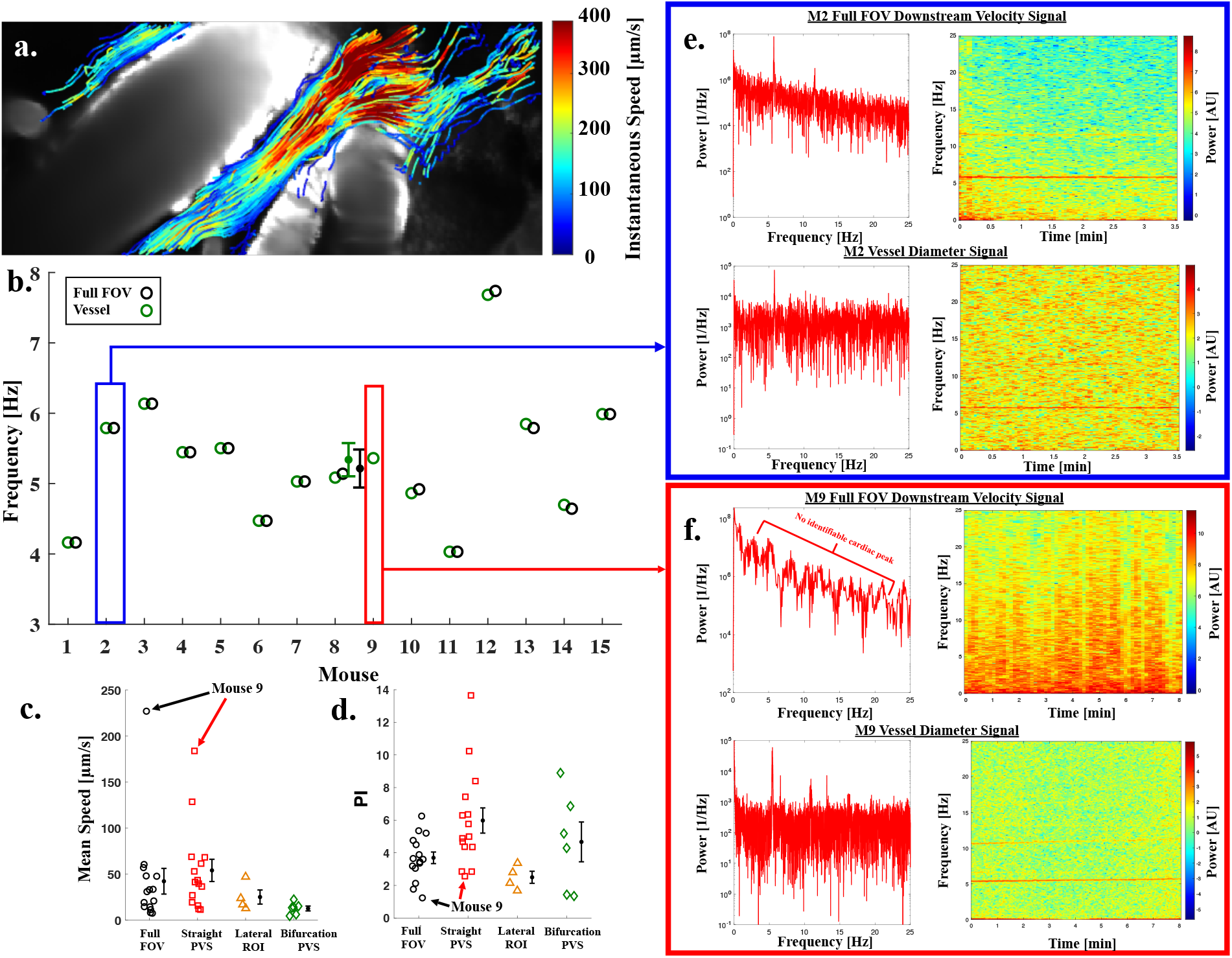
Analysis of an outlier mouse dataset exhibiting high mean flow speeds, low PI, and no identifiable *V*_downstream_ frequency peak in the power spectrum analysis. A localized constriction of the PVS within the FOV may contribute to fast but weakly pulsatile flow, highlighting the influence of anatomical heterogeneity on CSF transport dynamics. **(a)** Mouse 9 FOV with superimposed particle tracks plotted showing the instantaneous speed. Long solid-color streaks indicate the lack of pulsatility in the flow. **(b)** Principal frequency obtained from power spectrum analysis of the vessel diameter and *V*_downstream_ for all mice. **(c)** Mean flow speeds for mice and their respective ROIs showing that mouse 9 has a full FOV mean speed that is 8× that of the full FOV average. **(d)** PI of all mice and their respective ROIs showing mouse 9 with the lowest PI of any mouse in both the full FOV and straight PVS ROIs. **(e)** Power spectra (left) and spectrograms (right) for *V*_downstream_ (top) and vessel diameter measurements (bottom) from mouse 2, showing a clear peak that corresponds to the cardiac frequency. **(f)** Power spectra (left) and spectrograms (right) for *V*_downstream_ (top) and vessel diameter measurements (bottom) from mouse 9, showing no clear cardiac frequency peak for *V*_downstream_ despite a clear frequency peak in the vessel diameter measurements.

**Figure S3.**
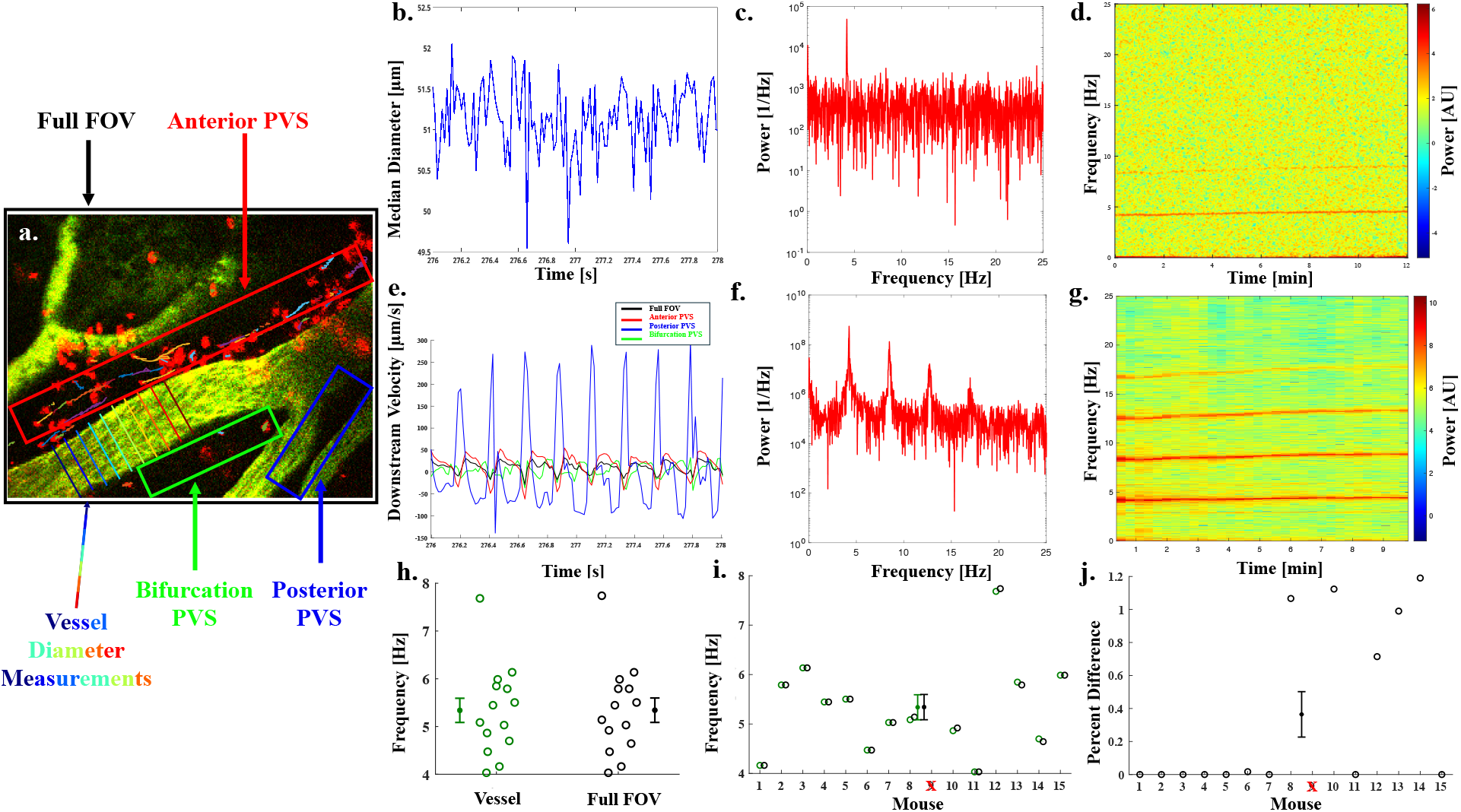
Power spectrum analysis of arterial diameter and *V*_downstream_ were measured and compared to verify that fluctuations in CSF velocity and arterial pulsations occur at the same frequency. **(a)** Vessel diameter lines and ROIs for each of the raw downstream velocity signals are depicted for a sample dataset. **(b)** Time snippet of raw arterial diameter measurements from vessel depicted in (a). **(c)** Power spectrum of the noisy signal from (b) shows a clear peak at the cardiacy frequency of 4.2 Hz. **(d)** Spectrogram showing a consistent vessel diameter frequency obtained from power spectrum analysis. This stable heart rate indicates appropriate anesthetic depth of the mouse during imaging. **(e)** Instantaneous *V*_downstream_ measurements in four different ROIs are shown during the same time snippet as (b). **(f)** Power spectrum of *V*_downstream_ from (e) shows the same frequency peak as that of (c) along with higher order harmonics. **(g)** Spectrogram of *V*_downstream_ for the full FOV ROI, showing a principal frequency consistent with that of the arterial diameter. **(h)** Principal frequency from power spectrum analysis of *V*_downstream_ for the full FOV ROI (black) and vessel diameter measurements (green) for *n* = 14 mice. The mean ± SEM for both is 5.4 ± 0.4 Hz. **(i)** Mouse by mouse comparison of the principal frequencies. **(j)** Percent difference between the *V*_downstream_ and vessel diameter principal frequencies plotted in (i), which yields an average difference of 0.6 ± 0.2%. In (h-j), only one recording per mouse (chosen based on the maximum number of particle tracks) was analyzed.

**Figure S4.**
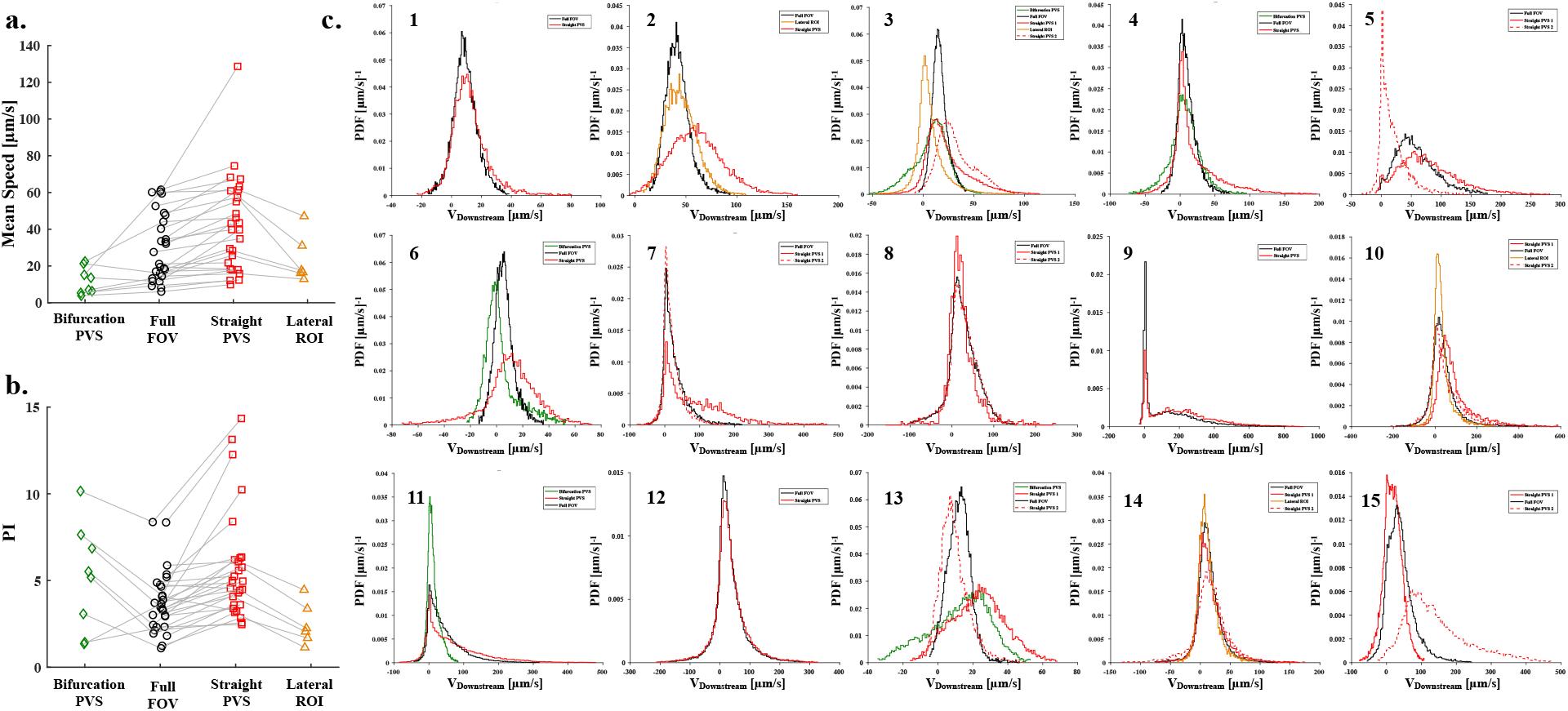
Characterization of mean flow speed, PI, and *V*_downstream_ for all mice. **(a-b)** Plots of (a) mean flow speed and (b) PI across all ROIs for each mouse. 28 recordings across *n* = 15 mice (1 to 4 recordings per mouse). Gray lines indicate measurements in the same recording. **(c)** Probability density functions of *V*_downstream_ for each mouse and its constitutive ROIs. Only one recording per mouse (chosen based on the maximum number of particle tracks) is plotted.

**Figure S5.**
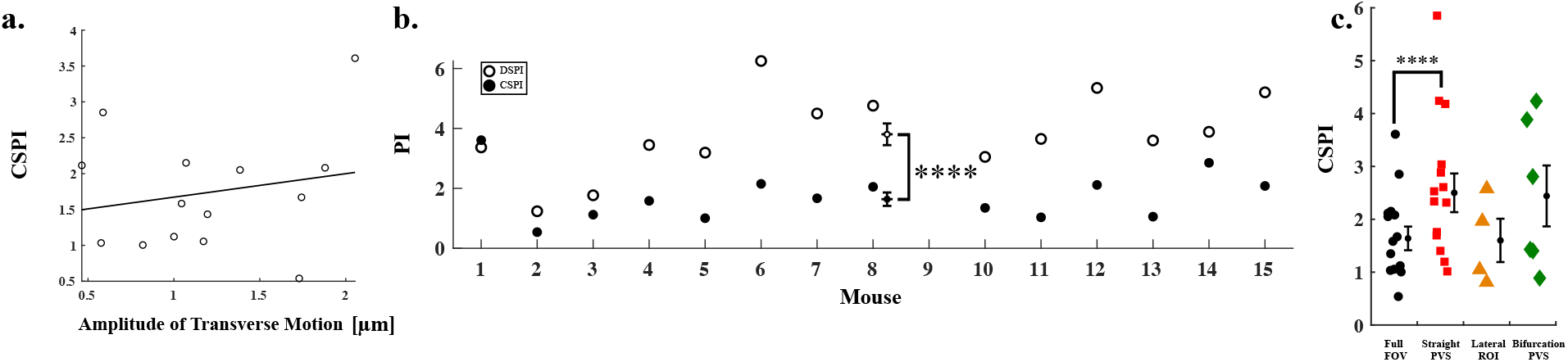
Cross-stream pulsatility index (CSPI), calculated from the cross-stream velocity component, exhibits different behavior from that of the downstream pulsatility index (DSPI) reported in the main text. **(a)** The CSPI shows weak correlation with the amplitude of transverse motion of the artery. **(b)** DSPI is larger than CSPI for 13 of 14 mice shown. Consistent with the main PI analyses, mouse 9 was excluded from statistical testing; DSPI was significantly larger than CSPI in the remaining mice (paired t-test, *p* = 2.04 × 10^−5^). Error bars near the center indicate the mean ± SEM. **(c)** CSPI is significantly larger for Straight PVS compared to Full FOV ROIs. Statistical comparisons among ROI types were performed using a linear mixed-effects model on log-transformed CSPI values, with ROI and frame rate as fixed effects and mouse and recording as random intercepts, followed by pairwise ROI comparisons using Holm correction across the six possible ROI comparisons (Full FOV vs Straight PVS, Holm-adjusted *p* = 1.35 × 10^−6^).

**Figure S6.**
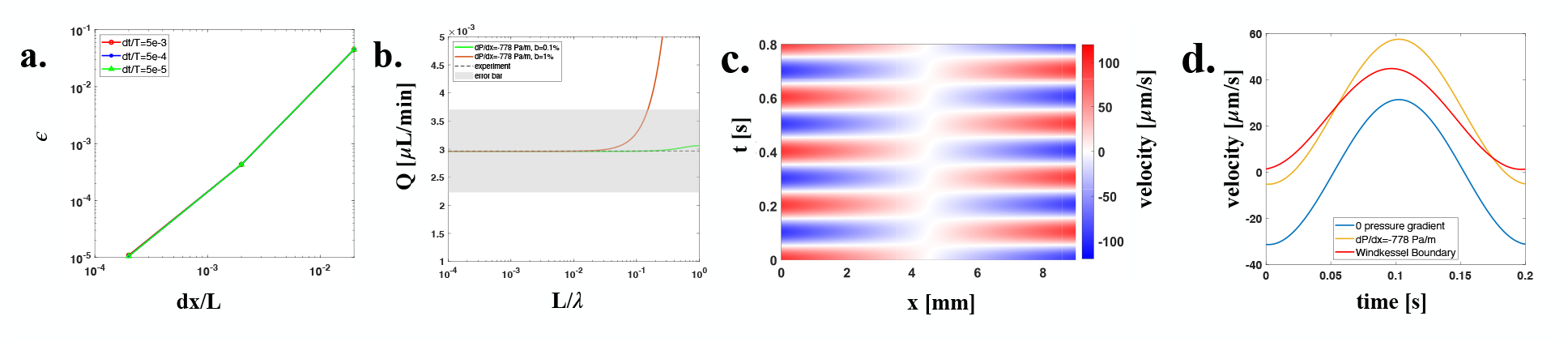
Numerical simulation verification and comparisons with experiment and across different models. **(a)** Verification of the numerical model via comparison against an analytical solution [31]. **(b)** Zoom-in view of comparison between the experimental and simulation results presented in Fig. 3b. **(c)** Spatiotemporal velocity color map of CSF flow in the PVS domain for the simulation without Windkessel boundary conditions. **(d)** Velocity waveform at *x* = 6 mm under different boundary conditions, indicated in the legend. For the Windkessel case (red), *R*_pial_ = 8.8 × 10^11^ Pa·s/m^3^, *R*_distal_ = 8.8 × 10^12^ Pa·s/m^3^, and *C* = 1.3 × 10^−15^ m^3^/Pa.

**Figure S7.**
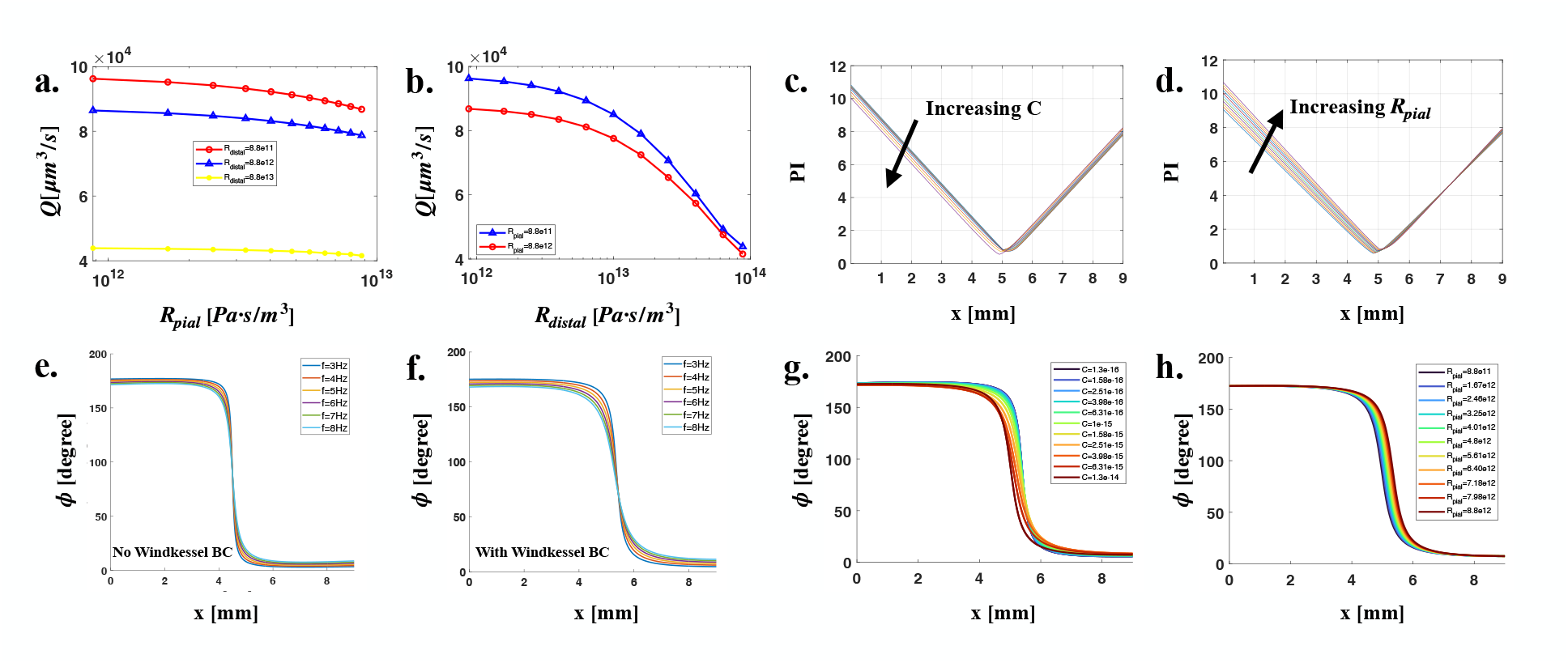
Additional results characterizing the finite volume 1D PVS model with Windkessel boundary conditions. **(a)** Variations of average volume flow rate with different pial and distal resistance values. *C* = 1.3 × 10^−15^ m^3^/Pa for all simulations. **(b)** Variations of average volume flow rate with different pial and distal resistance values (different projection view). *C* = 1.3 × 10^−15^ m^3^/Pa for all simulations. **(c)** Variations of pulsatility index, PI, along the domain with increasing compliance. *R*_pial_ = 8.8 × 10^12^ Pa s/m^3^ and *C* = 1.3 × 10^−15^ m^3^/Pa for all simulations. **(d)** Variations of PI along the domain with increasing pial resistance. *R*_distal_ = 8.8 × 10^12^ Pa s/m^3^ and *C* = 1.3 × 10^−15^ m^3^/Pa for all simulations. **(e-f)** Phase difference of arterial wall velocity and CSF flow velocity along the domain for different driving frequencies (e) without Windkessel boundary conditions and (f) with Windkessel boundary conditions. *R*_pial_ = 8.8 × 10^12^ Pa s/m^3^, *R*_distal_ = 8.8 × 10^12^ Pa s/m^3^ and *C* = 1.3 × 10^−15^ m^3^/Pa for all simulations. **(g)** Phase difference along the domain for different compliance values. *R*_pial_ = 8.8 × 10^12^ Pa s/m^3^ and *R*_distal_ = 8.8 1× 0^12^ Pa s/m^3^ for all simulations. **(h)** Phase difference along the domain for different pial resistance values. *R*_distal_ = 8.8 × 10^12^ Pa s/m^3^ and *C* = 1.3 × 10^−15^ m^3^/Pa for all simulations.

## A Numerical Methods

### A.1 One-dimensional finite volume method

In this section, we introduce a one-dimensional model to describe CSF flow within a single PVS segment. The numerical framework follows the approach described in [84] and is inspired by [85, 86]. The computational domain is simplified to an annular domain between two concentric cylinders, with a cross-sectional area that varies in space and time (Fig. 3a). Here, *r*_1_ represents the radius of the inner wall and *r*_2_ represents the radius of the outer wall. The governing equations are derived based on the finite volume method (FVM). In this model, the outer wall is assumed to be rigid, whereas the inner (arterial) wall is altered by a propagating peristaltic pulsation. The arterial pulsation can be modeled as a sinusoidal function dependent on both space (*x*) and time (*t*):

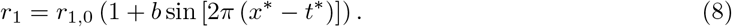

In this waveform, *r*_1,0_ represents the initial inner radius of the annulus, *b* is the amplitude of the wave, and *x*^∗^ and *t*^∗^ are the dimensionless space and time variables.

The CSF filling this annulus is assumed to be incompressible and Newtonian. In the FVM, the computational domain is discretized into a series of control volumes (CVs), arranged in a collocated grid (Fig. 3a). The east and west faces are shared with neighboring CVs, whereas the north and south faces coincide with the outer and inner walls of the annulus, respectively. The computational points are located at the center of each CV. For momentum conservation, each CV is treated as a hydraulic resistor based on Poiseuille flow in an annular geometry. The volume flow rate through the east or west faces of the *i*^*th*^ CV, *q*, is calculated using the hydraulic analog of Ohm’s law, involving local differences in pressure *P* and hydraulic resistance *R* (defined further below):

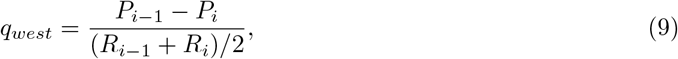

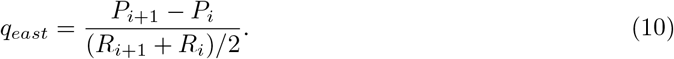

The pressure and flow in each CV is obtained from the change in fluid volume *V*, computed as the sum of fluxes through all faces. The east and west face contributions for the *i*^*th*^ CV are:

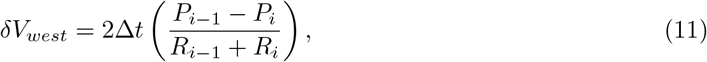

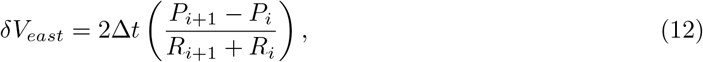

where Δ*t* is the time step. The volume change on the south face of the CV equals the volume displaced by the arterial wall *δV*_*artery*_, which varies in space and time due to arterial pulsation waves and is given by:

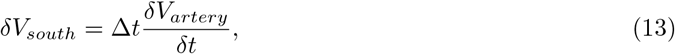

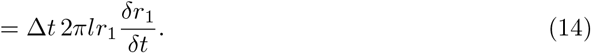

where *l* is the length of the CV. Since the outer wall is modeled as nearly rigid and impermeable, the volume change across the north face of the control volume is negligible and is approximated as zero:

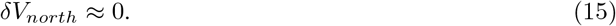

With the volumetric changes at each face of a CV known, the pressure, change in volume, and volume flow rate within each CV at time step *n* + 1 can be calculated as:

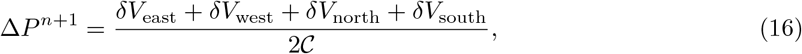

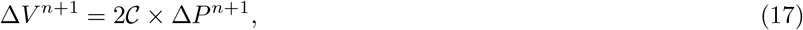

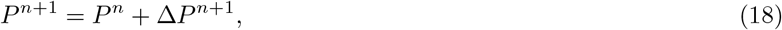

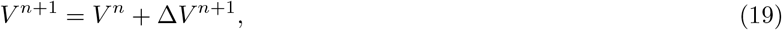

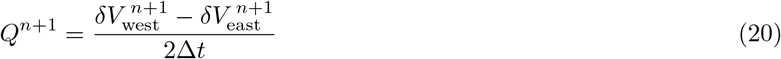

where *C* is the compliance of the brain tissue for the annular CV, computed as [84, 85]:

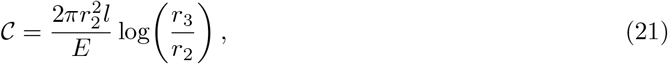

with Young’s modulus *E* = 10^12^ Pa, which is much larger than that of brain tissue, allowing the outer wall to be approximated as rigid [87]. Here, *r*_3_ is the typical length scale of the segment of parenchyma adjacent to the PVS, which we estimate based on a prior study of flow through penetrating PVSs [88].

A prior experimental study showed that the PVSs surrounding pial arteries are open (non-porous) spaces [89]. Accordingly, each pial PVS segment is represented as an open annulus, and its hydraulic resistance is computed under the assumption of a fully developed annular Poiseuille flow profile [90],

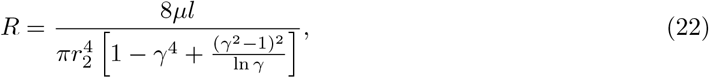

where *γ* = *r*_1_*/r*_2_.

Eqs. (16–20) for the *i*^*th*^ CV can be rearranged to obtain:

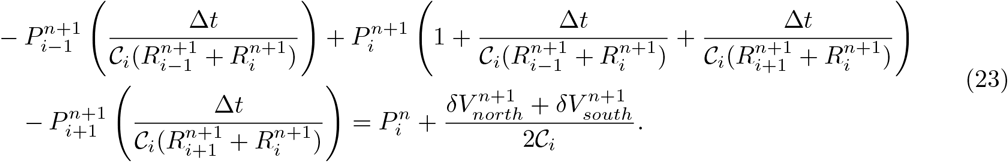

which is solved through direct inversion of a tridiagonal matrix under Dirichlet pressure boundary conditions. The corresponding left-hand side (LHS) and right-hand side (RHS) matrices are constructed as:

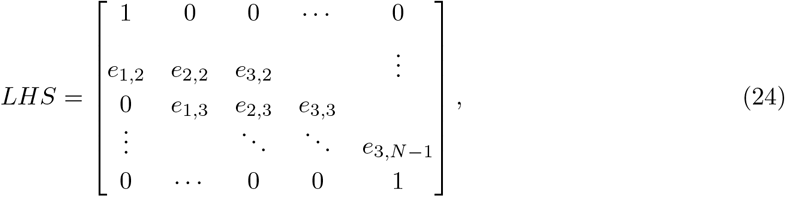

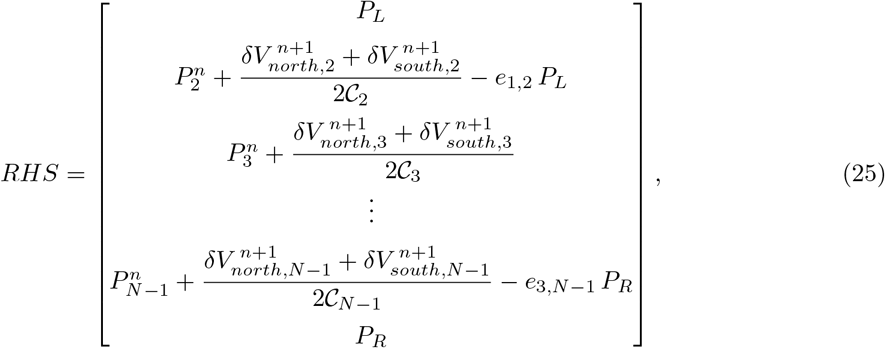

where *N* denotes the total number of CVs in the domain, *i* = 1, 2, …, *N*, and

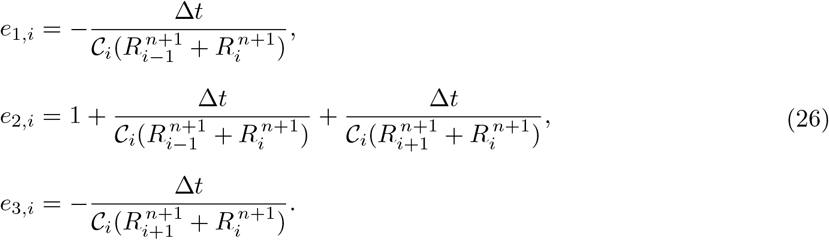

As previously described, the inner radius of the PVS varies according to a traveling sinusoidal wave given by Eq. (8). Based on the instantaneous inner radius, *R* is computed for each CV using Eq. (22). As the arterial pulsation propagates along the artery, the cross-sectional area of the PVS changes, thereby altering the local hydraulic resistance. Consequently, the flow resistance must be updated at each time step throughout the computational domain. After updating the pressure distribution, the volumetric flow rate is evaluated as:

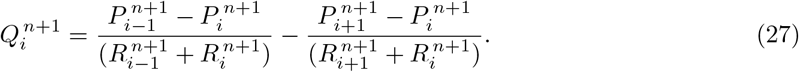

This finite volume model is a lumped-parameter formulation in which each CV is treated as a lumped hydraulic element characterized by its local resistance and compliance. This approach captures the essential physics of CSF flow while avoiding the computational complexity of directly solving the full Navier–Stokes equations. Compared with conventional CFD methods, this lumped-parameter FVM formulation offers several advantages: it is numerically stable and straightforward to implement, requires significantly less computational cost, and converges faster. In addition, the method is not subject to the strict Courant–Friedrichs–Lewy stability criterion that typically limits the time step size in explicit Navier–Stokes solvers, allowing for larger time steps and efficient long-term simulations of pulsatile CSF flow [84].

### A.2 Lumped parameter model for boundary conditions

Based on the finite volume framework described in the previous section, the FVM model initially employed Dirichlet boundary conditions with fixed inlet and outlet pressures. While this approach provides numerical stability, it does not accurately represent the physiological coupling between the PVS and the downstream channels in the network, which dynamically modulate the outlet pressure.

To capture these effects more realistically, a lumped parameter model, commonly referred to as a “Windkessel boundary condition,” is introduced. The Windkessel model treats the downstream vascular bed as a combination of resistive and capacitive elements, representing viscous losses and vessel/brain tissue compliance, respectively. By coupling the FVM domain with these lumped elements, the outlet boundary can respond dynamically to pulsatile flow, allowing more realistic modeling.

Inspired by the lumped-parameter framework reported by Ladrón-de-Guevara et al [54], we developed a three-element (RCR) Windkessel model to provide more physiologically realistic boundary conditions for the finite volume simulation. The model presented in that study employed a two-element Windkessel circuit, consisting of a distal resistance and a compliance element. Here, *C* represents the vascular and tissue compliance, which characterizes the capacity of the downstream flow channels to store fluid when pressure fluctuates. However, such a two-element configuration fails to reproduce the impedance characteristics of the proximal flow channels. Since our model neglects axial changes in the size of the PVSs, inclusion of a pial resistance element captures the resistive aspect of this neglected geometrical effect, improving the capacity for obtaining agreement with experimental measurements.

The governing equation of the RCR Windkessel boundary condition (located at the outlet of the modeled PVS domain) is:

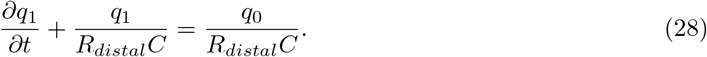

where *q*_0_ represents the flow rate at the PVS outlet (i.e., through *R*_pial_), while *q*_1_ denotes the flow rate through the *R*_distal_ (Fig. 3d). The numerical implementation of the Windkessel model is:

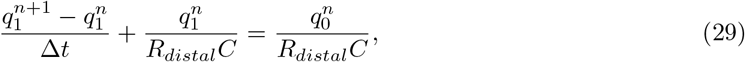

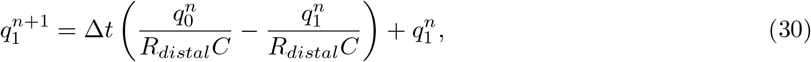

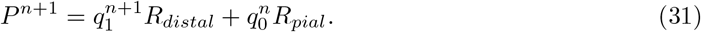

During each time step, the outlet pressure is dynamically updated according to the instantaneous flow and the state of the RCR circuit. This coupling enables the downstream boundary to respond realistically to pulsatile inflow conditions, rather than being constrained by a fixed or purely periodic pressure condition.

